# Bradyzoite subtypes rule the crossroads of *Toxoplasma* development

**DOI:** 10.1101/2025.08.23.671938

**Authors:** Arzu Ulu, Sandeep Srivastava, Nala Kachour, Michael W. White, Emma H. Wilson

**Affiliations:** Division of Biomedical Sciences, School of Medicine, University of California, Riverside, Riverside 92521, CA, United States

**Keywords:** *Toxoplasma gondii*, tissue cysts, surface antigens, chronic infection, bradyzoite replication

## Abstract

Reactivation of toxoplasmosis is a significant health threat to chronically infected individuals, especially those who are or become immunocompromised. An estimated one-third of the world population is infected with *Toxoplasma*, placing millions at risk. The *Toxoplasma* cyst is the foundation of disease with its ingestion leading to infection and its reactivation, from slow replicating bradyzoites to fast replicating tachyzoites, leading to cell lysis in tissues such as the brain. There are no treatments that prevent or eliminate cysts in part due to our poor understanding of the mechanisms that underlie cyst formation and recrudescence. In this study, we aimed to understand the biology of bradyzoites prior to recrudescence and the developmental pathways they initiate. We have discovered ME49EW cysts from infected mice harbor multiple bradyzoite subtypes that can be identified by their expression of distinct proteins. Sorting of these subtypes revealed they initiate distinct developmental pathways in animals and in primary astrocyte cell cultures. Single bradyzoite RNA sequencing indicates 5 major bradyzoite subtypes occur within these cysts. We further show that a crucial subtype comprising the majority of bradyzoites in chronically infected mice is absent from conventional in vitro models of bradyzoite development. Altogether this work establishes new foundational principles of *Toxoplasma* cyst development and reactivation that operate during the intermediate life cycle of *Toxoplasma*.

## Introduction

*Toxoplasma gondii* is a globally prevalent protozoan parasite, infecting nearly one-third of the human population. During chronic infection, the tissue cyst form is essential for both parasite survival and disease pathogenesis. In immunocompromised individuals, reactivation of these cysts, otherwise known as recrudescence, is the only known cause of acute, and sometimes fatal, toxoplasmosis [1, 2]. *Toxoplasma* reactivation is typically controlled by IFN-γ mediated immune responses [3]. However, with rising immunosuppression in the U.S. population, 6.6% of adults are immunocompromised, and with over a million cancer patients undergo chemotherapy annually, the risk of cyst reactivation becomes a growing public health concern [4]. Given its high global seroprevalence (up to 60% infections in some regions) [5, 6], and potential for severe disease, understanding bradyzoite biology and tissue cyst dynamics is critical.

*Toxoplasma* is a successful obligate intracellular parasite due to its broad host range and ability to form long-lived tissue cysts within neurons and muscle cells. Environmental transmission of *Toxoplasma* occurs via oocysts (containing sporozoites) that are shed by the feline definitive host, while the consumption of food contaminated with tissue cysts is the second route of *Toxoplasma* transmission. Bradyzoites within tissue cysts are the pivotal stage of the *Toxoplasma* life cycle. They serve as a reservoir of infection capable of differentiating directly into tachyzoites responsible for dissemination and cell lysis [7-9] or switching to the merozoite stage in the feline definitive host that leads to sexual reproduction in the cat gut [10-12]. Bradyzoites are also capable of direct replication, which is thought to be required for maintaining cyst burden and chronic infection [10, 11, 13-15]. There is great complexity in investigating this predominantly in vivo structure that contains slow growing parasites. Our recent methodology uses an ex vivo system of recrudescence relying on in vivo generated cysts [13]. This has allowed the characterization of the kinetics and phenotype of parasite growth following recrudescence that cannot be fully replicated using in vitro cell cultures and allows us to probe the heterogeneity and function of parasites prior and immediately after cyst reactivation.

In the *Toxoplasma* Type II ME49 strain, genes encoding SAG1-related surface proteins (SRS) includes 144 genes; 109 that match gene models and 35 incomplete pseudogenes [16, 17]. SRS genes are present on all ME49 chromosomes with clusters of SRS paralogs found on seven chromosomes. SRS antigens discovered 40 years ago remain an enigmatic class of *Toxoplasma* factors [18, 19]. There is evidence SRS antigens influence host cell adhesion [20], and it is postulated that immunodominant SRS antigens, such as SAG1, elicit immune responses against the tachyzoite stage so that bradyzoites, which lack SAG1 are spared [21]. Notwithstanding, the realization of purpose for a few SRS antigens, the function of the vast majority of SRS antigens remains unknown. Antibodies raised against developmentally regulated SRS antigens are valuable reagents in the study of *Toxoplasma* developmental pathways [19, 22]. In this respect, SAG1 antigen has served as a primary marker of the tachyzoite stage, while induction of SRS9 expression (i.e. SRS16B) indicates early bradyzoite development [13]. In this study we investigated the SRS22A protein that belongs to a cluster of 9 paralogs (SRS22A-I) and one pseudogene tandemly arrayed on the left end of chromosome VI. The SRS22 gene family are highly expressed in merozoites of the definitive life cycle [23, 24] with the exception of SRS22A mRNA, which is additionally expressed in bradyzoites [23].

In this study, we demonstrate SRS22A is a dominant bradyzoite surface protein expressed in ME49EW bradyzoites from mice but is not detected in conventional cell culture models of bradyzoite development, which now includes our ex vivo bradyzoite recrudescence model [13]. We determined that ME49EW tissue cysts from infected mice harbor at least two bradyzoite subtypes distinguishable by SRS22A expression that have different growth and developmental functions. Through the use of single bradyzoite mRNA-sequencing we confirm that in vivo bradyzoites differentially express SRS22A mRNA and provide evidence that there are additional bradyzoite subtypes likely present in the tissue cysts of chronically infected mice.

## Results

### Bradyzoites from chronically infected mice have unique surface antigen composition

It is generally understood that conventional alkaline-stress models of tachyzoite-to-bradyzoite differentiation do not faithfully reproduce the full breadth of native developmental processes. The strains used in these in vitro experiments often poorly form cysts in animals, the pH 8.2 media combined with CO2 starvation kills many parasites, and the gene expression changes induced by alkaline-stress are incomplete and developmentally confused [25]. The reliance on in vitro methods has contributed to significant gaps in our understanding of native bradyzoite developmental biology in the *Toxoplasma* intermediate life cycle. To begin addressing this problem, we sought to identify potential in vivo bradyzoite markers by comparing the mRNA expression of SRS genes in ME49EW bradyzoites from 30-day infected mice against in vitro ME49B7 tachyzoite and alkaline-stressed parasite mRNA expression (for full RNA-seq data see ref [13]). Seventeen SRS antigen mRNAs were expressed at least 10-fold higher in ME49EW bradyzoites from mice (Fig. 1A table). The mRNA encoding SRS9 (also designated as SRS16B) was highly expressed in ME49EW bradyzoites as well as alkaline-stressed in vitro bradyzoites validating the use of anti-SRS9 antibodies as an early bradyzoite marker in our studies. The gene encoding the SAG-related protein SRS22A was the top expressed SRS mRNA in ME49EW bradyzoites from in vivo cysts at >1000-fold higher levels than SRS22A mRNA levels in tachyzoites. The in vivo expression of SRS22A mRNA was also well above SRS22A mRNA levels from alkaline-stressed parasites (∼17 fold higher, Fig. 1A). We selected a peptide from the from the N-terminal region of the SRS22A protein coding sequence (*LRGNDGRSSRVIEKEAEVAK*; Fig. S1A) based on informatic prediction of immunogenicity and produced peptide-specific antibodies in rabbits. The SRS22A antisera successfully stained in vivo tissue cysts from mice (30-days post-infection), whereas there was no reactivity of the pre-immune serum control (Fig. S1B). Furthermore, Western blot analysis detected a strong, single band running slightly smaller than the 30kD molecular mass predicted in excysted bradyzoites obtained from 30-day infected mice with no signal detected in RH tachyzoites grown in vitro (Fig. 1B) indicating specificity for bradyzoites. It is common for SRS proteins to electrophorese faster than the predicted molecular mass in SDS-PAGE gels [26].

**Figure 1.**
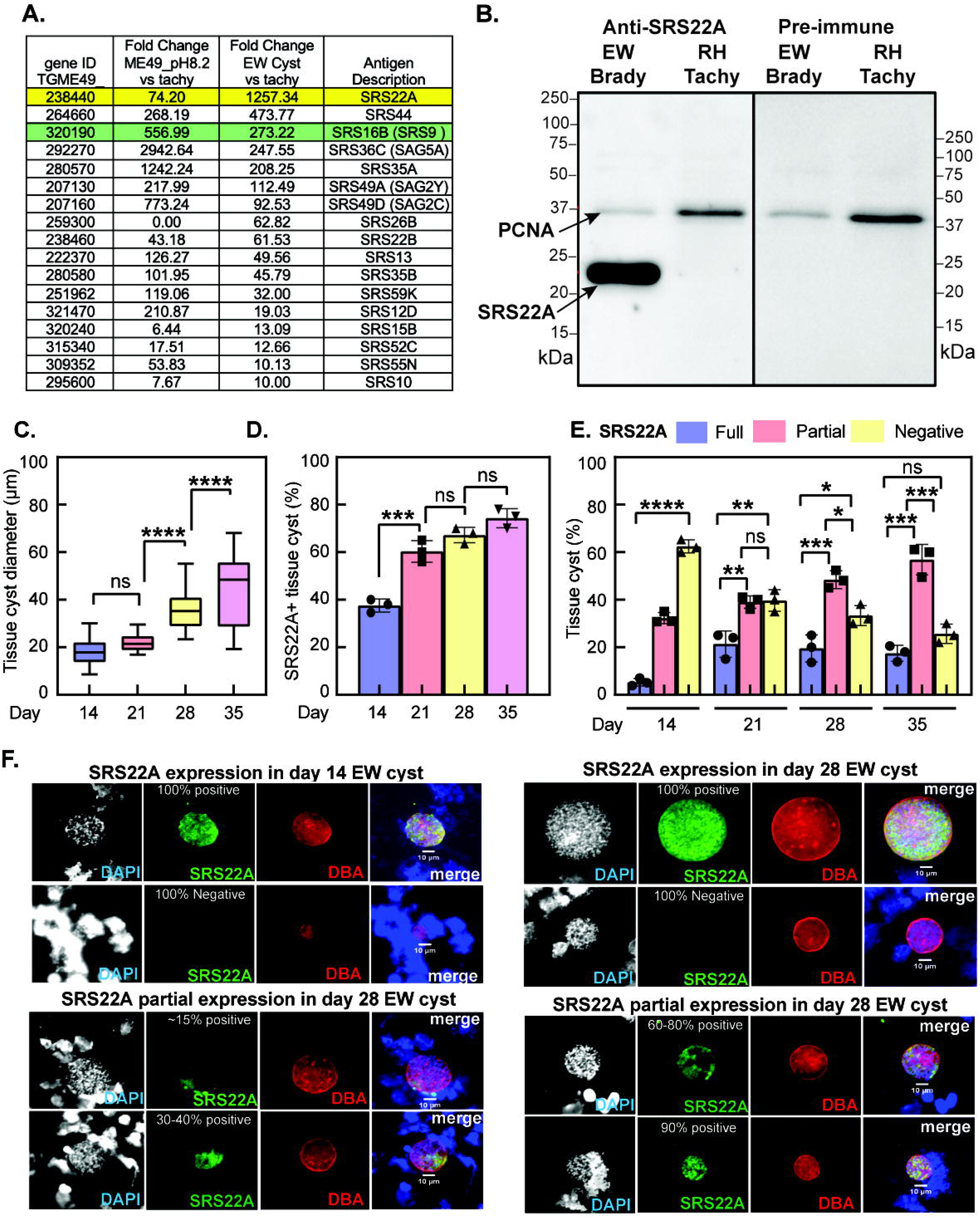
Identifying in vivo bradyzoite-specific SRS genes. **[A.]** Table of 17 SAG-related family genes (SRS) that have at least 10-fold higher expression in ME49EW in vivo bradyzoites (40-days post-infection) as compared to ME49B7 tachyzoites grown in vitro. Expression values of these SRS genes in ME49B7 tachyzoites subjected to 48 h of alkaline-stress (pH 8.2) were also included. SRS22A is the top expressed SRS gene in ME49EW in vivo bradyzoites, surpassing even SRS9 (SRS16B), a canonical bradyzoite marker of early bradyzoite development. SRS22A was chosen for peptide-based antibody development (see Fig. S1A). **[B.]** Western blot probed with rabbit anti-SRS22A antiserum detects a strong signal for SRS22A (∼22 kDa) in ME49EW bradyzoites (excysted from in vivo cysts) that is absent from RH strain tachyzoite controls. PCNA is included as a loading control. A pre-immune serum control is included in the right panel. **[C–E.]** Quantification of ME49EW tissue cysts in brain homogenates from CBA mice infected for 14-, 21-, 28- and 35- days: [C.] Tissue cysts in mouse brain increase in size over time. [D.] Percentage of tissue cysts harboring SRS22A+ bradyzoites, and [E.] Quantification of three SRS22A cyst phenotypes; 100% SRS22A+, partial SRS22A+ and 100% SRS22A- bradyzoites at the four timepoints. At least two brain homogenates were analyzed per timepoint, and each homogenate was counted 3x. **** = *P* < 0.0001, *** = *P* < 0.001, ** = *P* < 0.01, * = *P* < 0.05 **[F.]** Representative immunofluorescence images of tissue cysts at days 14- and 28-post-infection, showing SRS22A+ versus SRS22A- expression (homogenous or mixed). Scale bar = 10 μm. Brain homogenates were methanol-fixed and co-stained for SRS22A (green), DBA (red), and DAPI (blue) to indicate nuclei. Note that the interior regions of cysts that lack anti-SRS22A staining have numerous parasite nuclei. A decolorized DAPI image was included to better define the distribution of parasite nuclei in each cyst.

Utilizing the SRS22A antiserum, we determined the kinetics and cyst distribution of SRS22A expression in tissue cysts from infected brain homogenates at 14-, 21-, 28-, and 35-days post-infection (Figs. 1C-F). Although there are likely complex patterns of cyst growth, rupture and reformation over the course of infection, average cyst size generally increases over time [12, 13], which is consistent with our results here (Fig. 1C). The proportion of cysts expressing SRS22A is generally correlated with increasing cyst size (Fig. 1D). By the last timepoint (Day 35), >75% of tissue cysts showed some expression of SRS22A. Over the course of infection, we estimated that cysts exhibited three SRS22A bradyzoite configurations; near 100% positive or negative or mixtures of SRS22A+/SRS22A- bradyzoites (see Fig. 1F for examples). The proportion of ∼100% positive SRS22A cysts increased from Day 14 to Day 21 post-infection, and this level of expression was maintained at ∼20% of total cysts thereafter (Fig. 1E). By contrast, the proportion of cysts with mixtures of SRS22A- and SRS22A+ bradyzoites continued to increase over time reaching >50% of the population by Day 35. A process of cyst maturation has been postulated to explain the differences in cysts generated by in vitro alkaline-stress models (considered to be early cysts) and the putative mature cysts recovered from >30-day infected mice. By probing for SRS22A, (an indicator of in vivo cysts) further cyst heterogeneity is revealed and indicates there is no single "mature" tissue cyst in the native intermediate life cycle, but that cyst heterogeneity exists throughout the course of *Toxoplasma* infection in the brain.

### SRS22A is exclusively expressed by in vivo generated cysts

SRS22A protein expression in tissue cysts from mice validated the SRS mRNA expression results for this antigen (Fig. 1A). We next determined whether SRS22A was expressed in bradyzoites that develop in cell culture models. SRS22A mRNA expression in ME49EW cysts from mice is substantially higher than in cysts generated under alkaline-stress treatment (Fig. 1A). To determine if SRS22A protein expression differentiates in vivo from invitro bradyzoites, we used ME49B7 tachyzoites [27] to infect HFF monolayers, and following a 4 h invasion period, shifted half the cultures to pH 7.4 media and the other half to pH 8.2 bradyzoite-media. We co-stained infected HFF cells with two pairs of antibodies; the tachyzoite-specific SAG1 monoclonal antibody with the bradyzoite-specific anti-SRS9 or anti-SRS22A rabbit antisera. Parasite vacuole size (2- 64 size vacuoles) and relative SRS antigen expression was assessed at 48 h after the shift to pH 8.2 media (Fig. 2A). ME49B7 tachyzoites grown in pH 7.4-media actively replicated and were positive for SAG1 antigen expression (Fig. 2A, graph plots). In pH 8.2-media, ME49B7 tachyzoites grew much slower as evidenced by the smaller range of vacuole sizes. Tachyzoite vacuoles (SAG1+ only) were 41% of the total parasite population whereas 59% of vacuoles co-stained for SAG1+ and SRS9+ (D+ vacuoles) or expressed only SRS9+ antigen, which were considered to be bradyzoite vacuoles. No parasites in either pH 7.4- or pH 8.2-media expressed detectable SRS22A antigen.

**Figure 2.**
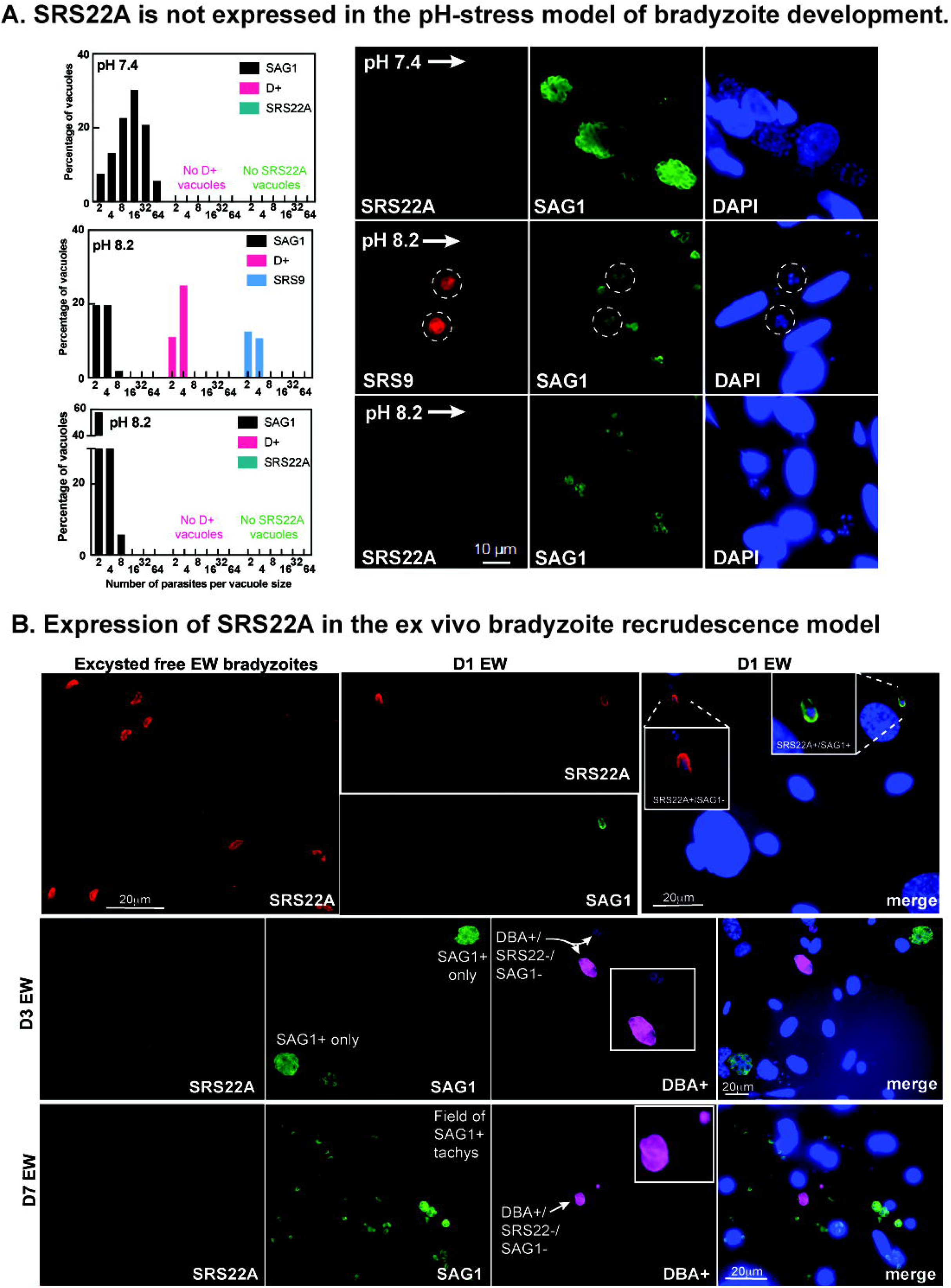
Cell culture models of bradyzoite development fail to express SRS22A. **[A.]** (right images) SRS22A expression was not detected in any parasite vacuole irrespective of the pH condition used. The presence of SRS9+ vacuoles that were the result of tachyzoite-to-bradyzoite development in pH 8.2 media (48 h) confirm early bradyzoite differentiation in these cultures. The infected HFF cell cultures were fixed and co-stained for SRS9 (bradyzoites, red) and SAG1 (tachyzoites, green), or separately for SRS22A (red) and SAG1 (green), which was required because SRS9 and SRS22A antisera were raised in rabbits. DAPI was included to visualize nuclei. Scale bar = 10 μm. (left graphs) Quantification of vacuole size for SAG1+ only, double positive SAG1+/SRS9+, and SRS9+ only vacuoles demonstrated cultivation in pH 8.2 media caused overall slower parasite growth, which is known to be associated with the alkaline-stress bradyzoite differentiation model. **[B.]** Immunofluorescence images showing SRS22A staining in ME49EW excysted, free bradyzoites and ex vivo bradyzoite-infected primary astrocyte cultures at days 1-, 3-, and 7- post-infection. Free bradyzoites and infected-astrocyte cultures were co-stained for SRS22A (red), SAG1 (tachyzoite-specific, green), and DBA (cyst wall-specific, magenta). Notably, SRS22A staining diminishes very quickly (within the 1st 24 h) in ex vivo bradyzoite-infected astrocytes, and thus, at day 3 (D3 EW) and day 7 (D7 EW) post-infection no parasites stained positive for SRS22A. Note that SRS22A was also not expressed or re-expressed in bradyzoite vacuoles where cyst wall formation was active (DBA+). Scale bar = 20 μm.

In our ex vivo recrudescence model, in vivo ME49EW bradyzoites are used to infect primary astrocytes cultivated under low oxygen (5%) conditions [13]. As expected, SRS22A antisera stained excysted ME49EW bradyzoites from in vivo cysts as well as single bradyzoites that had invaded astrocytes (Fig. 2B, free and Day 1 bradyzoites). The majority of vacuoles in Day 1-infected astrocytes contain single (pre-replication) bradyzoites reflecting time needed for dormant bradyzoites to reawaken [13]. In advance of the first parasite divisions, we detected switches in SRS antigen composition indicating some changes in developmental transcription (e.g. SRS22A) occur before replication commences in full. The two-representative single bradyzoite vacuoles shown here (Fig. 2B, D1 EW) both express SRS22A, while only one vacuole co-stained brightly for SAG1. In the ex vivo recrudescence model [13] replicating bradyzoite vacuoles occur alongside tachyzoite vacuoles (SAG1+) as visualized by DBA positive cyst walls (Fig. 2B). Representative examples of developmentally mixed populations of bradyzoites and tachyzoites (DBA+ versus SAG1+ staining) in Day 3- and 7-infected astrocytes are shown (Fig. 2A; D3 EW and D7 EW image panels). Importantly, SRS22A expression was not observed in these mixed populations (Fig. 2B) in either the fast-replicating tachyzoite period at Day 3 (D3 EW) or the post-growth shift tachyzoite/bradyzoite population at Day 7 (D7 EW). Similar to the alkaline-stress model no expression of SRS22A antigen was detected in any parasite vacuole beyond Day 1 in the infected astrocyte cultures.

### Bradyzoite subtypes initiate different developmental pathways

The discovery of SRS22A+ bradyzoites in tissue cysts of infected mice, but not in cell culture models of cyst development, raised questions about the function of both SRS22A- and SRS22A+ bradyzoite subtypes. The finding that cysts from chronically infected mice had mixed SRS22A (+ and -) bradyzoite populations (Fig. 1E) gave equal importance to investigating SRS22A- bradyzoites. Therefore, to determine the developmental function of these bradyzoite subtypes, we sorted SRS22A+ and SRS22A- bradyzoites from brain tissue cysts by FACS (Fig. 3, see also Fig. S2 controls). ME49EW cysts were isolated from 40-day infected mice using a Percoll gradient and bradyzoites excysted (D0 population) as previously described [13]. The purified bradyzoites were then stained using anti-SRS22A antibody. Identification and cell sorting of parasite populations revealed a distinct SRS22A+ expressing population representing approximately 60- 80% of all Day 0 bradyzoites. To determine any differential SRS22A-associated growth patterns, sorted populations were used to infect primary astrocyte cultures (Fig. 3A). Population growth and SRS antigen expression patterns (SAG1 versus SRS9) were determined during the maximal ex vivo growth stage at Day 5 post-infection. The infected cultures revealed clear differences in growth rates. SRS22A+_seeded cultures expanded 3.1-fold by Day 5, whereas parasite populations from SRS22A-_seeded cultures only reached replacement levels (1.1-fold). Remarkably, the end-stage parasites (tachyzoites or bradyzoites) produced by these cultures were also different. On average over 80% of sorted SRS22A expressing bradyzoites converted to replicating tachyzoites (SAG1+ only) with very few bradyzoite vacuoles (2.3%, SRS9+ only). In contrast, bradyzoites that were originally SRS22A-led to the opposite developmental profile and replicated as bradyzoites expressing SRS9 and forming DBA positive cysts (Fig. 3A,B). Here, bradyzoites (30% of vacuoles, SRS9+ only) were the principal end-stage produced with few tachyzoite vacuoles present (3.7% vacuoles SAG1+ only). Tissue cyst wall assembly was only detected in the SRS9+/SAG1- vacuoles (Fig. 3B). We used a restrictive criterion for calling end-stages, and thus, in both SRS22A infections there were intermediate parasites (see examples Fig. S1C) co-expressing varying levels of SAG1 and SRS9 antigens (pairwise dull to bright staining; 16% in SRS22A+, >60% in SRS22A-) and made up the remaining proportion of parasites. The lower growth rate of SRS22A-seeded cultures likely contributed to the slower developmental progression of these populations, although it was clear that the transient double positive parasites rarely resolved to the SAG1+ only tachyzoite stage as they did in abundance in the SRS22A+ bradyzoite-infected astrocytes.

**Figure 3.**
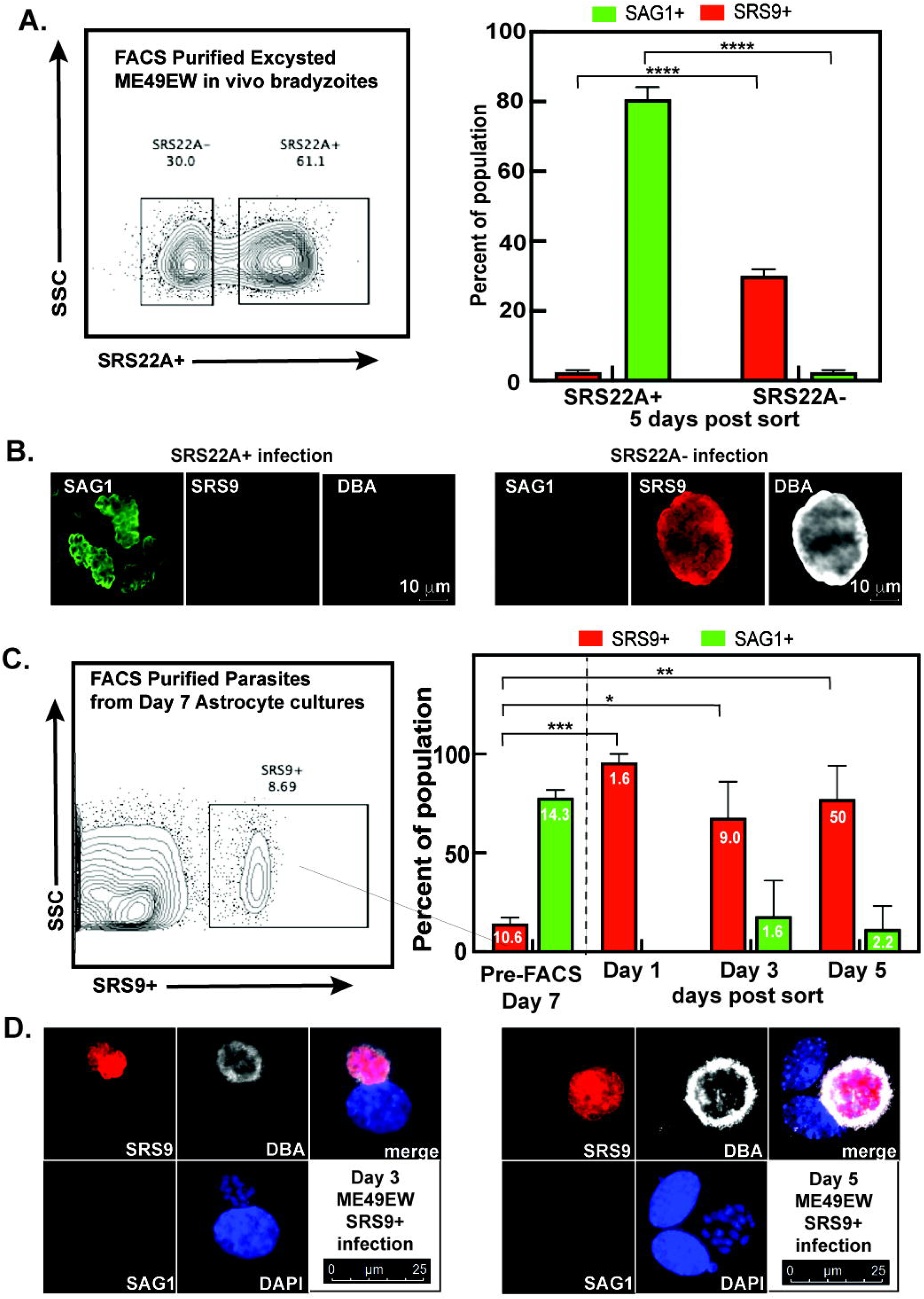
Developmental fate of SRS22A bradyzoite subtypes. **[A.]** Excysted ME49EW bradyzoites from CBA mice brain tissue (40-day infection) were subjected to FACS purification in order to purify SRS22A bradyzoite subtypes (see Fig. S2A for FACS control plots). Flow cytometric analysis revealed the pre-sorted ME49EW bradyzoite populations from these mice were 63% SRS22A+ and 36% SRS22A-. Primary mouse astrocyte monolayers grown on microscope cover slips were inoculated with purified SRS22A+ or SRS22A- bradyzoites (500 parasites/cover slip). At 5-days post-infection the parasites were fixed and co- stained with anti-SRS9 and anti-SAG1 antibodies followed by DAPI staining. The total number of parasites was determined microscopically for each coverslip (3.1-fold increase for SRS22A+, 1.1-fold increase for SRS22A-). Three replicate counts of end-stage SAG1+ only and SRS9+ only vacuoles were also performed and graphed here. **[B.]** We observed cyst wall formation (as detected by DBA+ staining) only in SRS9+ bradyzoite vacuoles that were the major product of the SRS22A- infections as shown by the representative images. Scale bar = 10 μm **[C.]** SRS9+ only (SAG1-) parasites from ME49EW Day 7-recrudescing populations (ex vivo bradyzoite recrudescence,[13]) were also purified by flow cytometry and used to infect primary astrocytes. As noted in Figure 2A, Day-7 ME49EW parasites do not express SRS22A. Triplicate counts of vacuole size (included column numbers indicate average vacuole size) and SAG1+ only or SRS9+ only antigen expression collected at 1-, 3-, and 5-days post-infection were graphed. No SAG1+ only vacuoles were detected in Day 1 populations. Parasite vacuoles expressing variable levels of both SRS antigens were also present at 4-14% of the total vacuoles at each timepoint (not graphed). The average vacuole size and end-stage fractions in the original pre-sort Day 7- population were also indicated by column number or by bar graph, respectively. **[D.]** Representative images of day-3 and -5 post-infection vacuoles in astrocytes infected with purified SRS9+ bradyzoites are shown. Scale bar = 25 μm. **** = *P* < 0.0001, *** = *P* < 0.001, ** = *P* < 0.01, * = *P* < 0.05.

The preference of SRS22A- bradyzoites to initiate further bradyzoite replication suggests two major aspects of bradyzoite-cyst biology: firstly, that there is heterogeneity of bradyzoites within a cyst and secondly that these subpopulations have preferential developmental trajectories. Our previous work revealed heterogeneity of bradyzoites during ex vivo recrudescence with a proportion of parasites (∼20%) replicating as SRS9+ bradyzoites (brady-brady replication) in addition to the canonical brady-tachy pathway associated with cell lysis and pathology during cyst rupture [13]. To determine if these subpopulations of bradyzoites also represent a stable functional phenotype, SRS9+ bradyzoites were sorted from recrudescing populations in astrocytes and evaluated for average vacuole size and SAG1 versus SRS9 antigen expression at Day 1-, 3- and 5-days post-infection (Day 7 parasites, Fig. 3C). As we demonstrated in Fig. 2, Day 7-bradyzoites spontaneously assemble tissue cyst walls (DBA+) but do not express SRS22A, and are therefore, SRS9+/SRS22A-. Pre-sort, SRS9+ parasites (undergoing brady-brady replication) represented ∼8% of the population (Fig. 3C) and as expected sorted parasites retained the SRS9+ only pattern 24 h after inoculation (Fig. 3C, Day 1). However, these parasites continued to express only SRS9+ over the course of the next 5 days and did not express SAG1. It was apparent that these bradyzoites retained their ability to replicate, evidenced by the increase in SRS9+ only vacuole size (Fig. 3C, column numbers). After 5 days of sorted parasites being in culture ∼75% of vacuoles remained brady-brady replicating parasites successfully making cyst wall proteins (Fig. 3D). A proportion of parasites (∼20%) did convert to SAG1+ only vacuoles, however, these parasites were not growing well (average vacuole size of <2). It is notable that tachyzoites in the pre-FACS Day 7-population grew better with an average vacuole size of 14.3 likely because the parentage of these parasites was the fast-growing tachyzoite [13]. Altogether these data demonstrate that in vivo SRS22A- as well as in vitro SRS9+/SRS22A- bradyzoites are primary initiators of direct bradyzoite expansion, while in vivo SRS22A+ bradyzoites are responsible for recrudescence into the fast-growing tachyzoite stage.

### Infection patency requires SRS22A+ bradyzoites

To evaluate whether FACS-purified SRS22A+ and SRS22A- bradyzoite populations are functionally and developmentally competent, we infected CBA mice with each bradyzoite subtype. Our goal was to assess parasite burden at two key time points: acute phase (5-days post-infection) and chronic phase (28-days post-infection). At the acute time point (5-days post-infection), we observed a significantly greater number of SAG1+ tachyzoites in peritoneal exudate cells (PECs) from SRS22A+ infections compared to SRS22A- infections (*P* < 0.05; Fig. 4A). Quantification of parasite-vacuoles revealed more SAG1+ vacuoles in the SRS22A+ infections (Fig. 4A). Additionally, we assessed the ability of SRS22A+ vs SRS22A- parasites to disseminate and cause systemic infection by measuring parasite burden in the liver, spleen and lung of infected mice (Fig. 4B). While all tissues showed a trend towards increased parasite burden in the SRS22A+ infections relative to the SRS22A- infections, the differences did not reach statistical significance. However, at 28-days post-infection, quantification of brain cysts revealed that mice infected with SRS22A+ bradyzoites developed significantly more cysts than those infected with SRS22A- bradyzoites (*P* < 0.05; Fig. 4C). This is consistent with the robust dissemination of fast-growing tachyzoites [13] that SRS22A+ bradyzoites produce.

**Figure 4.**
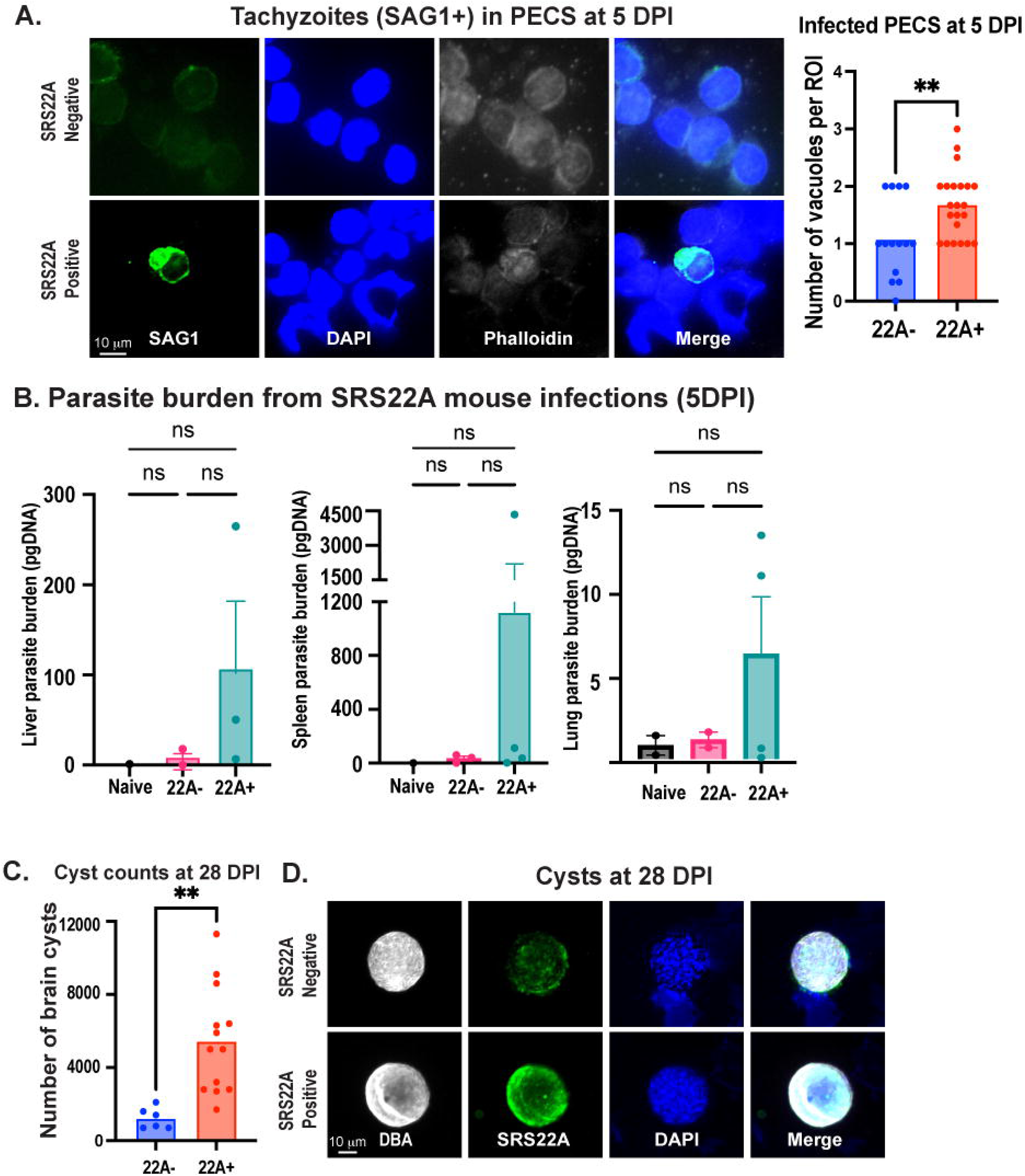
In vivo studies of FACS purified SRS22A+ and SRS22A- bradyzoites. **[A.]** Representative images of peritoneal exudate cells (PECs) at 5-days post-infection (DPI). PECs were collected from each mouse by injecting intraperitoneally with 5 ml cold PBS, and 100,000 cells per sample were deposited onto microscope slides using cytospin funnels. Right panel: quantification of the number of SAG1⁺ parasite vacuoles per region of interest (ROI). Data are mean ± SEM; *n* = 3 mice per group. **[B.]** Parasite burden in the liver, spleen and lung was assessed by qPCR targeting the *Toxoplasma B1* gene. Data are mean ± SEM. Liver parasite burden: *n* = 1 (naïve), *n* = 2 (22A⁻), *n* = 3 (22A⁺); spleen and lung parasite burden: *n* = 1 (naïve), *n* = 3 (22A⁻), *n* = 4 (22A⁺). **** = *P* < 0.0001, *** = *P* < 0.001, ** = *P* < 0.01, * = *P* < 0.05. **[C.]** Mice were infected intraperitoneally with 10,000 FACS-purified SRS22A+ or SRS22A- bradyzoites. At 28-days DPI, brain cysts were enumerated from whole brain homogenates. Homogenization was performed sequentially using 18G, 20G, and 22G needles. Data are presented as mean ± SEM; *n* = 6 mice for 22A⁻ and *n* = 13 for 22A⁺ infections. Representative immunofluorescence images of cysts collected at 28-days post-infection from the infections described in (A). Brain homogenates were smeared onto microscope slides, fixed in methanol, and stained with anti-SRS22A antiserum. and *n* = 24 cysts (SRS22A⁺ infections).

### Single cell transcriptome analysis of in vivo bradyzoites provides additional evidence of multiple bradyzoite types

The above data using only 3 antigens, demonstrate heterogeneity of bradyzoites that predicts the functional development of parasites. To further our understanding of the underlying basis for this heterogeneity, we analyzed the transcriptome of single, excysted ME49EW bradyzoites from 40-day infected mice (Figs. 5, 6, S3, and S4). Individual bradyzoites from two brain tissue cyst harvests collected more than 6 months apart (Fig. S3A, r=0.965) were captured using the 10x Chromium Single Cell Gel Bead kit and 3’- RNA-seq libraries sequenced. Sequencing of these cRNA libraries produced ∼22 billion reads with >80% of reads mapping to the *Toxoplasma* genome and a total of 4,190 bradyzoites passed quality controls (Fig. S3B). The scRNA-seq data were embedded using Uniform Manifold Approximation and Project (UMAP) and parasite transcriptome intersection determined using the Seurat package (R version 4.4.2 and Seurat 5.1.0). UMAP projections of bradyzoites from each cyst harvest generated matching parasite clusters (Groups A-E) irrespective of the bradyzoite sample analyzed (Fig. S3C). The merged bradyzoite data (Fig. 5A, UMAP projection) yielded a range of ∼1,200-3,300 mRNAs in each bradyzoite Group (average mRNA values, <0.05 p-val, Database S1) with 374 genes expressed in all Groups (Fig. 5B, Venn diagram). The largest number of unique transcripts were identified in Groups A, B, D and E, while Group C, which interfaces with Groups A, B, and D, had the fewest (<75) unique transcripts. *Toxoplasma* is unusual for the number of mRNAs that are cell cycle regulated with ∼40% of *Toxoplasma* encoded genes showing periodic transcription profiles with peak mRNA levels in G1 or in S/M phases [28]. Previous studies of the tachyzoite cell cycle transcriptome also uncovered a potential relationship between bradyzoite-specific gene expression and parasites in the S/M cell cycle phases [28]. The scRNA sequencing of in vivo bradyzoites appears to confirm this connection with Group A-E gene expression (Fig. 5C, table) exhibiting nearly double (except Group D) the expected S/M cell cycle transcriptional influence (17% expected), whereas the proportion of G1 transcripts were similar to the predicted fractional value (∼23% expected). Nevertheless, bradyzoite Group A-E transcriptomes did not phenocopy the synchronous cell cycle contours as we uncovered when scRNA-sequencing ME49EW tachyzoites from our ex vivo recrudescence model [13]. Furthermore, analysis of the top principal components indicate that most of the variance between parasites is not driven by cell cycle stage or canonical bradyzoite markers (BAG1, LDH2, ENO1) (Fig. S3E). This contrasts with previous studies using alkaline stress, where these factors contributed more significantly to parasite variation [25, 29]. It is understood that cell cycle markers are at best indirect methods of measuring cell proliferation. Thus, there is no protein marker that is able to accurately quantify in situ *Toxoplasma* parasite replication without additional evidence. DNA synthesis and/or content methods are the most direct, reliable, and accurate methods to measure cell proliferation. Our previous flow cytometric analysis of in vivo bradyzoite genomic DNA content [13] revealed that ∼2% of these bradyzoites possessed a 1N+ DNA content (∼50% of the cell cycle). Group E bradyzoites had the largest number of S/M transcripts (745 mRNAs) and these bradyzoites displayed uniformly higher expression of key growth gene mRNAs (Fig. S3D; see also Fig S4C, A vs E heat map). Therefore, it is likely Group E bradyzoites are the small replicating fraction of otherwise dormant populations of bradyzoites in the tissue cysts of chronically infected mice [13].

**Figure 5.**
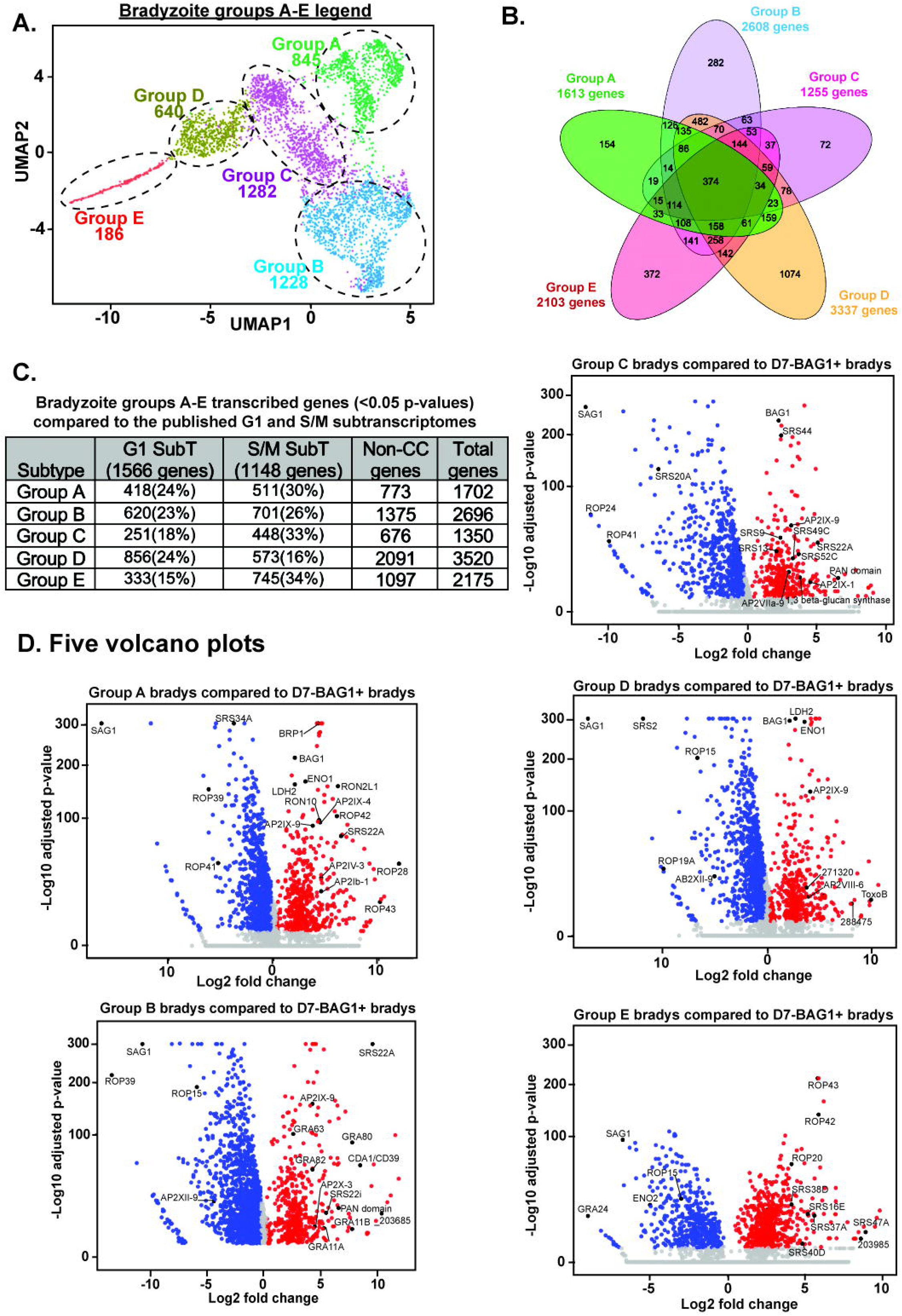
Single cell RNA-sequencing of in vivo bradyzoites. **[A.]** UMAP projection of individual in vivo bradyzoites from 30-day infected CBA mice shows five potential subtypes; Groups A, B, C, D, and E with the number of bradyzoites analyzed indicated below each group label. **[B.]** A Venn Diagram of significant gene expression (P <0.05) from bradyzoite Groups A-E was generated. This analysis shows all common, partially shared and unique genes associated with each Group. **[C.]** Table indicates the total number of genes in Groups A-E that are significantly expressed (Database S1, P <0.05) and the breakdown of shared expression with the published G1 and S/M sub-transcriptomes [14, 28, 37, 51] as well as the number of non-cell cycle genes. **[D.]** Volcano plots for each Group A-E were generated in comparison to BAG1+ bradyzoites in day 7 ex vivo cultures of infected astrocytes. Grey indicates genes with p>0.05, red indicated significantly upregulated genes (P < 0.05) and blue indicates downregulated genes (P < 0.05). Up to 15 upregulated and 5 downregulated genes are indicated on each volcano plot.

**Figure 6.**
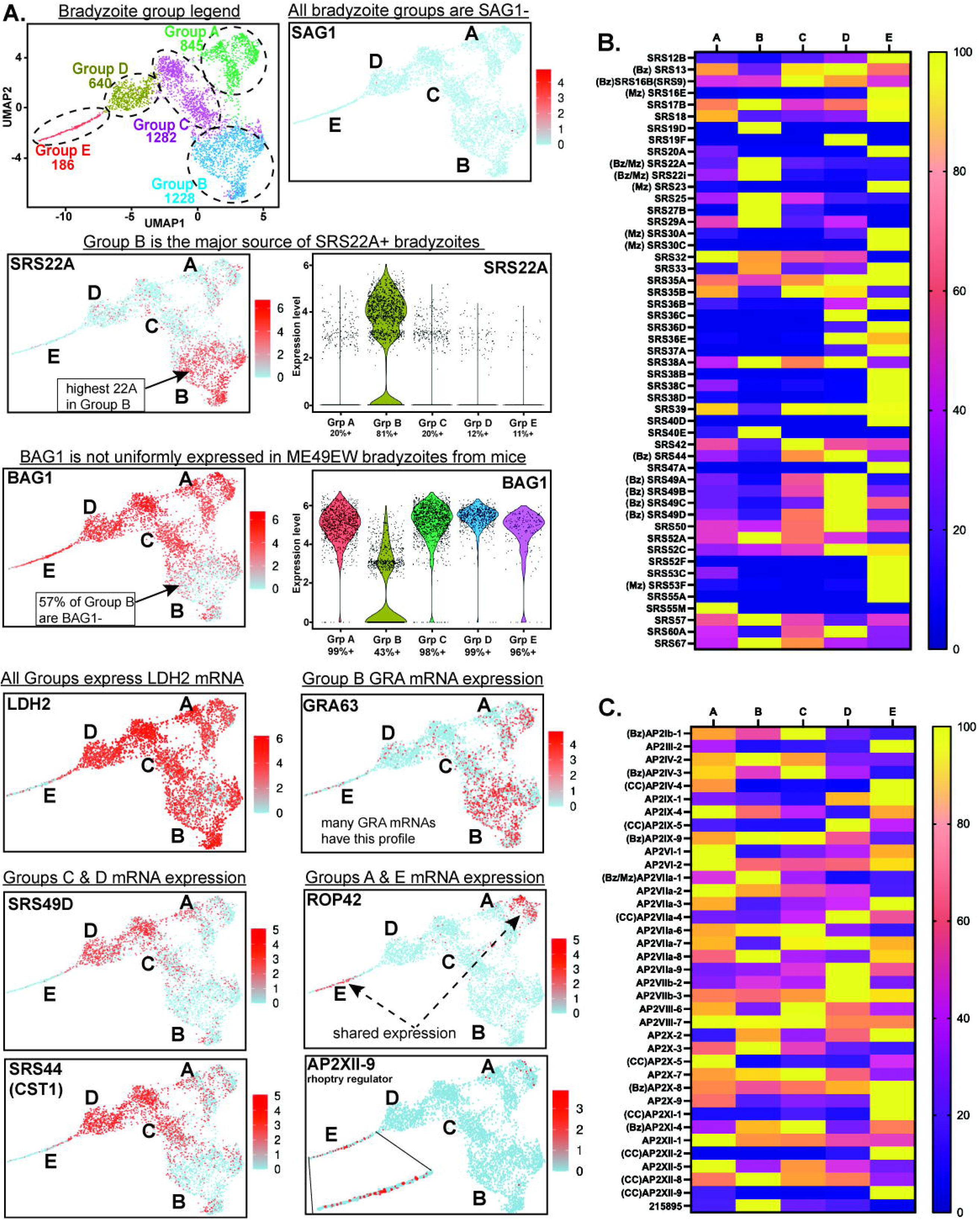
Gene expression profiles of five bradyzoite subtypes. **[A.]** (**top row**) The left UMAP image of individual in vivo bradyzoites that clustered in Groups A-E is included as a reference. UMAP image of SAG1 expression shading is on the right. (**second row**) UMAP projection and violin plot of SRS22A expression highlights expression in Group B bradyzoites (81% of all bradyzoites expressing SRS22A). (**third row**) UMAP projection and violin plot of BAG1 shows non-uniform expression among the clusters with Group B having the least expression of BAG1 mRNA. (**fourth row**) UMAP image of LDH2 and GRA 63 expression. Violin plot of LDH2 expression and a heat map of dense granule gene expression is shown in Fig. S4. (**left UMAP images in rows five and six**). Groups C and D shared expression of SRS49D and SRS44 (CST1). (**right UMAP images rows five and six**) Shared expression of ROP42 and AP2XII-9 in Groups A and E. Group E expression image of AP2XII-9 is magnified. **[B and C.]** SRS and ApiAP2 heat maps of normalized mRNA expression in bradyzoites from mice. SRS and ApiAP2 genes lacking expression in Groups A-E were excluded. Bz: bradyzoite, Mz: merozoite and CC: cell cycle specific genes.

To determine the possible functional differences between each cluster of bradyzoites found in cysts, differentially regulated transcripts in Groups A-E were compared to the transcriptome of Day-7 BAG1+ ME49EW bradyzoites. These bradyzoites spontaneously form in primary astrocytes after ex vivo recrudescence [13] and represent a replicating, cyst forming and developmentally competent population of bradyzoites in which to compare our originating bradyzoite cyst clusters. A combined UMAP projection of the in vivo versus ex vivo bradyzoites demonstrated clear separation of the in vivo bradyzoite Groups (A-E) from the distinct cluster of D7-BAG1+ bradyzoites from astrocytes (Fig. S3F). Hundreds of up-and down-regulated mRNAs were detected in Group A-E bradyzoites (Fig. S3F, table) compared to D7-BAG1+ bradyzoites (log2 fold-change results, Database S2) with many of the highest up-regulated mRNAs encoding recognized canonical bradyzoite protein markers (e.g. BAG1, LDH2, ENO1), selective rhoptry and GRA proteins, ApiAP2 factors, and SRS proteins as indicated (Fig. 5C volcano plots; see also Database S2). Considering the significant functions of these proteins in *Toxoplasma* biology, we also compared their expression among Groups A-E. The average mRNA values (Database S1) of AP2 factors and SRS protein stood out as exhibiting distinct patterns for each bradyzoite Group with AP2IX-9 the highest expressed ApiAP2 factor in bradyzoite Groups A-D (Fig. S4A). The mRNA encoding tachyzoite-specific antigen, SAG1, was the most significantly down-regulated transcript in Group A-E bradyzoites (Fig. 5D, labeled in volcano plots), which is visualized by the absence of SAG1 mRNA shading of the in vivo bradyzoite UMAP projection (Fig. 6A). The presence of SAG1 on the surface of parasites is a distinguishing marker bradyzoites that develop in cell culture models from the in vivo bradyzoites of infected mice, respectively [13].

In the experiments above, we established the patterns of in vivo bradyzoite SRS22A antigen expression and determined the distinct developmental functions of SRS22A+ and SRS22A- bradyzoite subtypes (Fig. 3). Thus, the expression profile of SRS22A mRNA in bradyzoite Groups A-E was of considerable interest. Indeed, SRS22A transcript expression was found highly focused in Group B bradyzoites accounting for 81% of the bradyzoites exhibiting the highest UMAP SRS22A estimates of mRNA levels (Fig. 6A). Consistent with the lack of SRS22A protein expression in cell culture-formed bradyzoites (Fig. 2), we did not detect SRS22A mRNA in D7-BAG1+ bradyzoites (Fig. S3G). Several other features of Group B bradyzoite transcription were notable. (1) The small heat shock protein, BAG1 was the first bradyzoite-specific marker characterized [30-32] and for more that 30 years BAG1 has been considered to be a universal marker of bradyzoites. It was therefore unexpected that over half of Group B bradyzoites lacked detectable expression of BAG1 mRNA (Fig. 6A). By contrast, Group B bradyzoites as well as bradyzoite Groups A, C & D and to a lesser extent Group E bradyzoites expressed high levels of LDH2 mRNA (Fig. 6A; Fig. S4A) supporting LDH2, a lactate dehydrogenase, as a more reliable marker for in vivo bradyzoites. (2) The SRS22 family of surface proteins are primarily merozoite-specific [23, 24] and Group B bradyzoites expressed two SRS22 members, SRS22A as mentioned and also SRS22i mRNA (Fig. 6B, SRS heat map). The analysis of the Group B bradyzoites lacking BAG1-identified two other SRS22 family members, SRS22E and SRS22G, although the expression of these SRS22 family members were lower than SRS22A and SRS22i (Database S1). (3) It may be related that Group B bradyzoites also had the highest levels of 24 dense granule protein (GRA) mRNAs including bradyzoite-specific GRA46 and GRA63 (Fig. 6A, GRA63) as well as merozoite GRA proteins GRA11A and GRA11B and GRA80-82 (Fig. S4A,B). (4) Finally, Group B bradyzoites expressed significantly higher levels (e.g. 21-fold higher Group B vs D, Database S1) of merozoite transcription factor, AP2VIIa-1 (Fig. 6C)[24]. Altogether, these features may indicate Group B bradyzoites are poised to initiate merozoite development in the cat. Further study will be required to understand whether Group B bradyzoites or another in vivo bradyzoite Group are the primary drivers of bradyzoite-to-merozoite development.

In addition to the distinct signature of Group B SRS22A expression, the in vivo bradyzoite Groups A-E displayed other distinctive SRS mRNA profiles (Fig. 6B, heat map). Ten SRS protein mRNAs had peak levels in Group E bradyzoites (Fig. 6B heat map, see also Database S1), including four merozoite-specific SRS proteins, SRS16E, 23, 30C, 53F and one SRS mRNA encoding a tachyzoite-specific protein SRS20A. The SRS mRNA profiles of bradyzoite Groups C and D were partially shared reflecting the nearly 70% overlap of Group C transcripts with the transcriptome of Group D bradyzoites. In particular, the expression of the SRS49 gene family mRNAs (SRS49A-D) was a distinct feature of bradyzoite Groups C and D (Fig. 6A,B). Group D bradyzoites had the highest proportion of unique mRNA expression (Fig. 5B, 1074 mRNAs) and also one of the lowest levels of SRS22A expression indicating Group D is likely a major source of SRS22A- bradyzoites. Intriguingly, the highest levels of SRS9 mRNA expression were in Group C and D, which correlates with the SRS22A- bradyzoite phenotype (see Fig. 3). The results above demonstrated SRS22A- bradyzoites initiate bradyzoite-to-bradyzoite development and it was SRS22A- bradyzoite infections that were responsible for spontaneous cyst wall formation (Fig 3 and [13]). Consistent with these findings, Group C and D bradyzoites expressed the highest levels of SRS mRNAs that encode major cyst wall proteins, 75% of bradyzoites expressing SRS44 (CST1) (Fig. 6A) and 62% of bradyzoites expressing SRS13 (Fig. S4A) were in Groups C and D.

Group A bradyzoites express the 3rd highest number of unique transcripts (Fig. 5B, 172 mRNAs) that includes many hypothetical proteins and increased expression of mRNAs encoding selective rhoptry and many RON mRNAs (e.g. Fig. 6A, Fig. S4C). This characteristic was partially shared with Group E bradyzoites and includes elevated expression of BRP1 and other bradyzoite-specific rhoptry proteins notably ROPs 28, 30, 42, 43 and RON2L1 mRNAs (Fig. S4A) [23, 24, 33] and the Group A-enhanced ROP kinase (TGME49_308093, Fig. S4). Increased expression of mRNAs encoding rhoptry proteins (e.g. >50-fold for some ROP mRNAs), which are S/M transcripts, was not a consequence of cell cycle phase synchrony as other S/M transcripts [34] were not elevated in Group A bradyzoites (Fig. S4C, A vs E heat map). By contrast, the increased expression of rhoptry related mRNAs did correlate with the increase of other S/M transcripts in Group E (Fig. S4C), consistent with the active replication of these bradyzoites. The mechanism responsible for increased expression of rhoptry and RON mRNAs in Groups A and E may involve AP2XII-9 which was also elevated in Group A and E (Fig. 6A). Recent studies have demonstrated a primary role for AP2XII-9 in activating rhoptry and RON gene transcription [35, 36]. We do not fully understand the biological role for increased rhoptry gene transcription in Group A bradyzoites. Our previous cell cycle studies of rhoptry proteins in tachyzoites [37] demonstrated the transcription of this class of mRNAs was uniquely sensitive to intercellular egress and invasion, which might indicate Group A bradyzoites were preparing to egress prior to tissue cyst harvest.

These scRNAseq data support the SRS22 and SRS9 findings that cysts contain distinct subsets of bradyzoites that lead to predictable patterns of development. These 5 populations separate based, not on cell cycle, but signatures that include characteristic proteins involved in parasite invasion and developmental regulation suggesting some of these populations are precursors to merozoites, tachyzoites and renewal of the bradyzoite population.

## Discussion

In these studies, we describe three major advances in our understanding of bradyzoite biology. Firstly, the expression of the SAG related protein, SRS22A, occurs specifically in bradyzoites formed in vivo and defines a population that transforms to the fast replicating, disseminating tachyzoite. Secondly, the alternative population found following recrudescence, brady-brady replicating parasites, represents a SRS22A- population that is stable and maintains the propensity to replicate as a bradyzoite (Fig. 7 model). Lastly, using scRNAseq of excysted bradyzoites, we observed further bradyzoite heterogeneity represented by 5 main groups of parasites identifiable by their expression of AP2 transcription factors and SRS proteins.

**Figure 7.**
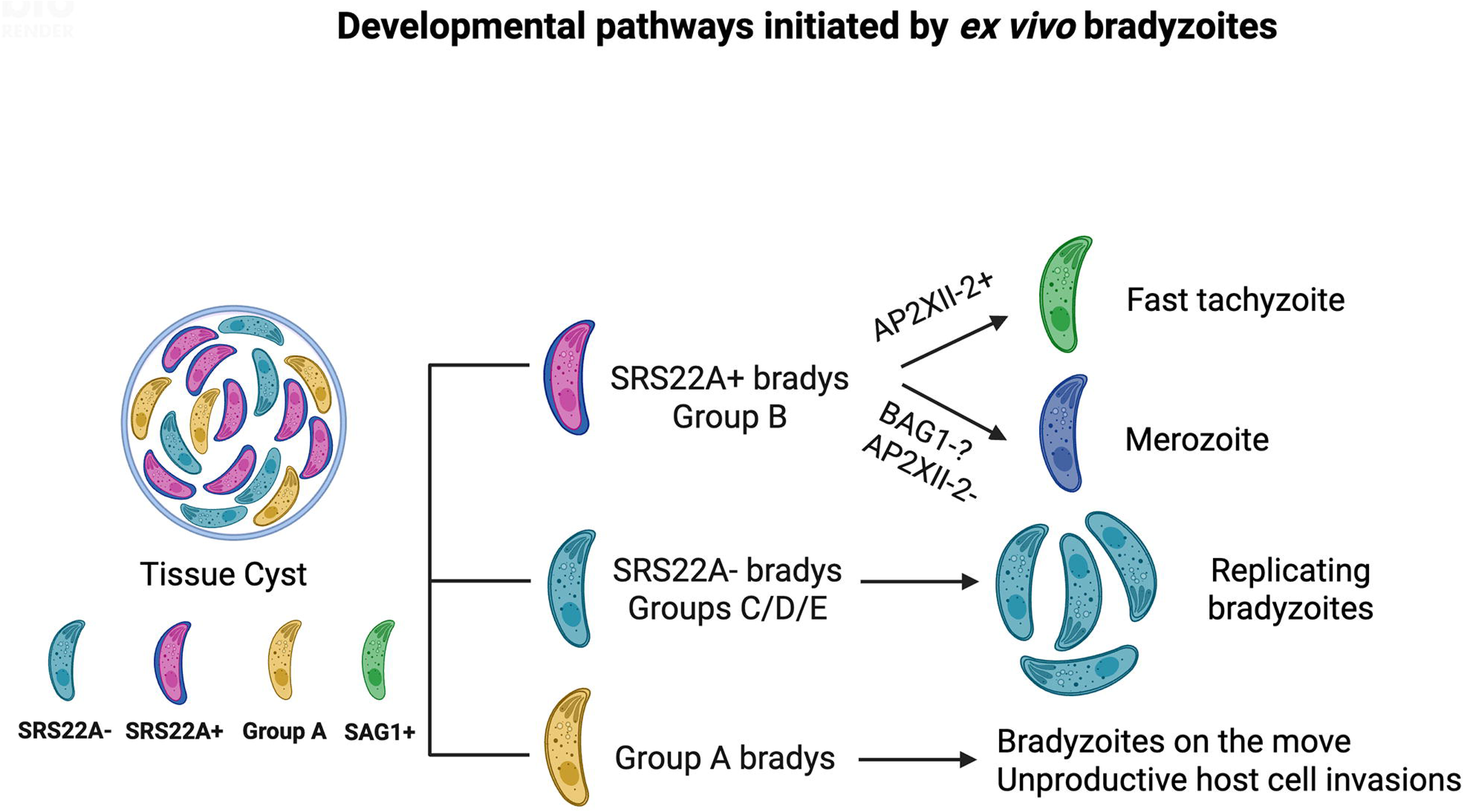
Model illustrating the non-linear life cycle and multifunctional nature of bradyzoites within heterogeneous tissue cysts. Our findings reveal that bradyzoites both positive and negative for SRS22A antigen can coexist within the same tissue cyst. SRS22A expression distinguishes two bradyzoite subtypes that initiate different developmental pathways: SRS22A+ bradyzoite infection initiates bradyzoite recrudescence into fast-replicating tachyzoites, while infections with SRS22A− bradyzoites initiates bradyzoite-to-bradyzoite replication. A large fraction of SRS22A+ bradyzoites are BAG1-, and it is possible these bradyzoites may initiate the definitive life cycle in feline host, which will require alterations in AP2 factor expression. Rhoptry related gene expression is triggered by egress and invasion events. We have found that a group of in vivo bradyzoites (Group A) possesses this pattern of gene expression, which may help explain the large number of documented parasite invasion events that do not lead to productive infections of host cells.

Our previous studies of sporozoites from cats and bradyzoites from mice demonstrated a common developmental sequence follows sporozoite or bradyzoite ex vivo infections of host cell cultures [13, 14, 38, 39]. This begins with a switch to a fast-growing tachyzoite stage followed by a near synchronous shift to slower tachyzoite growth 6-7 days later, after which bradyzoites begin to develop from the slower growing tachyzoite populations. A later reversion back to the fast-growing tachyzoite has never been observed in these cultures and the fast-tachyzoite only forms in ex vivo sporozoite and bradyzoite infections. It is not understood why fast-growing tachyzoites that are competent to form cysts in mice [13] have a restricted developmental context nor has any molecular signature emerged that provides an explanation for how they form. In this study, we advanced our understanding of the ingredients needed to produce natural fast-growing tachyzoites. Purification of ME49EW SRS22A+ bradyzoites from infected mice by excystation and flow cytometry and subsequent infection of astrocytes clearly established that the parentage of fast-growing tachyzoites is the SRS22A+ bradyzoite (Fig. 3). We show here that no in vitro model of bradyzoite development we tested was capable of forming the SRS22A+ bradyzoites (Fig. 2), and this explains why fast-tachyzoites have not been observed in *Toxoplasma* in vitro experiments. Our recent studies of ME49EW fast-growing tachyzoites determined they possess enhanced efficiency for dissemination in mice [13], which offers a possible explanation for why this stage might be needed in the intermediate host life cycle.

The most common view of parasite development in the *Toxoplasma* intermediate life cycle is a linear construction [40]. It is thought that tachyzoites differentiate in a limited stepwise fashion into a single end-stage, mature bradyzoite. As noted above, bradyzoite- or sporozoite-initiated development is assumed to only differ by the extra step of switching back to the tachyzoite before starting the linear pathway. Our recent studies of bradyzoite recrudescence [13] discovered replicating bradyzoites in tissue cysts forming spontaneously alongside the more prevalent tachyzoite vacuoles in astrocytes [13], confounding the simple linear view of bradyzoite development. We now understand that the origin of these distinct replicating stages was not a single parent, but instead two distinct parents present in our ex vivo inoculum. The SRS22A- bradyzoite subtype was the parent of bradyzoite-to-bradyzoite tissue cysts and the SRS22A+ bradyzoite subtype was responsible for the dominant bradyzoite-to-tachyzoite recrudescence. Thus, bradyzoite development is not linear but rather the result of multiple active pathways progressing in parallel. What is surprising is that different pathways are initiated by distinct bradyzoite subtypes raising the question of how many bradyzoite subtypes there are given that bradyzoites initiate at least three different pathways: brady-brady, brady-tachy (and the reverse) and brady-merozoite (Fig. 7 model). Importantly, based on our results following SRS22A+ and SRS22A- bradyzoite development (Fig. 1), we found that subtypes are formed early, and tissue cysts are heterogenic with some exhibiting uniform expression and others only partially positive for SRS22A. This heterogeneity is established within two weeks of infection and the proportion of cysts uniformly expressing SRS22A stabilizes at around 20% of the total cyst population with the rest being made up of cysts containing SRS22A+ and SRS22A- bradyzoites. This points to a desired characteristic of biology to have diversity at all levels and specifically in *Toxoplasma* infection of having diversity of bradyzoites within and between cysts. Moreover, the SRS22A subtypes (+ or -) are stable, can be purified and the developmental preferences are active in cell culture and in mice. This developmental arrangement would afford individual tissue cysts the ability to successfully transmit the *Toxoplasma* infection to a wide range of animal hosts. Based on our scRNA-sequencing of in vivo bradyzoites, we estimate there could be up to 4-5 bradyzoite subtypes within a single tissue cyst. We do not fully understand how bradyzoite subtypes are regulated, although it is likely that epigenetics plays a major role, and it is clear that in vivo bradyzoite groups A-E express distinct ApiAP2 factors (Fig. 6C). Supporting these concepts, recent studies of ME49EW mutant bradyzoites demonstrated that genetic knockout of ApiAP2 transcription factor, AP2IX-9 (TGME49_306620), or the bradyzoite cyclin, TgCYC5 (TGME49_293280), disrupted the balance of SRS22A+ versus SRS22A- cyst subtypes, which in turn caused shifts in bradyzoite recrudescence commiserate with the altered SRS22A phenotype in each bradyzoite mutant [41].

The heterogeneity of SRS22A+ bradyzoites (Fig. 3) was confirmed at the mRNA level with Group B bradyzoites the major source of SRS22A+ expression (Fig. 6). Group B bradyzoites have a unique and unexpected phenotype with >50% of these bradyzoites lacking expression of BAG1. Group B bradyzoites also had the lowest expression of SRS9 and CST1 (SRS44) and are the drivers of differentiation into fast-replicating tachyzoites (Fig. 3 and 6). Group B bradyzoites had the highest expression of SRS22A mRNA as well as mRNAs encoding SRS22i, SRS22E and SRS22G, and dense granule proteins GRA11, and GRA80-82, which are all merozoite-specific proteins. Turning off BAG1 expression may signal a priming event that pre-stages SRS22A+ Group B bradyzoites to recrudescence into the tachyzoite stage and also into the merozoite stage provided a suitable host environment. Our first encounter with priming of developmental gene expression was studying *Eimeria bovis* sporozoites and merozoites indicating this phenomenon is an ancient property of Coccidian parasites [42]. Both tachyzoites and merozoites lack BAG1 expression and they are the major replication stages for their respective life cycles. Consequently, they have shared metabolic needs and derive energy through similar biochemical mechanisms [23]. Supporting the premise of a common parent are recent studies that demonstrate altering a single AP2 factor [43, 44] can switch parasite replication from endodyogeny of the tachyzoite to endopolygeny of the merozoite [45]. Further studies to investigate the functional breadth of Group B bradyzoites are underway.

We showed above that SRS22A- bradyzoites are the primary initiator of bradyzoite replication and it is in these vacuoles that spontaneous cyst wall formation occurs in ex vivo infections of astrocytes ([13], Fig. 3). RNA sequencing established that bradyzoite Groups C and D are likely the major source of SRS22A− bradyzoites (Fig. 6) and it is these bradyzoite groups that we observed the highest levels of SRS9 mRNA and mRNAs encoding cyst wall proteins CST1 and SRS13, consistent with their trajectory toward tissue cyst formation. Like BAG1, SRS9 does not appear to be a canonical marker of in vivo bradyzoite development and the preference of SRS9 expression in SRS22A- bradyzoites from mice explains why purified SRS9+/SRS22A- bradyzoites from astrocytes phenocopied the developmental course of in vivo SRS22A- bradyzoite recrudescence (Fig. 3B). Previous data [13] initiated the hypothesis that slower replicating bradyzoites post-recrudescence (brady-brady) were responsible for long term cyst maintenance in chronically infected mice and are not the major drivers of tissue dissemination during the acute stage of infection. The infection of mice with purified SRS22A+ and SRS22A- bradyzoites supports this concept. In mice infected with purified SRS22A+ bradyzoites, we observed more SAG1+ parasites in PECS at the acute phase of the infection suggesting an increase in dissemination, which was later reflected in the chronic phase with increased cyst burden in this group as opposed to infections initiated with purified SRS22A- bradyzoites (Fig. 7).

The function of Group A bradyzoites is less clear. Our RNA sequencing results established that Group A bradyzoites upregulate many mRNAs encoding rhoptry effector proteins (Fig. S4C). In earlier studies, we found the transcription of this class of genes to be uniquely sensitive to parasite egress and host cell invasion [37]. There is substantial evidence that *Toxoplasma* invades many host cells without establishing a parasitophorous vacuole or producing a tissue cyst [46, 47]. It is therefore possible that Group A bradyzoites represent parasites that are in preparation of egress from the tissue cyst and primed for host cell invasion. This phenomenon has been observed with bradyzoites indicating that bradyzoite motility is just as effective as tachyzoites for invasion [48-50]. Intriguingly, we have found that the deletion of bradyzoite-specific cyclin, TgCYC5, may cause an increase in these bradyzoite dissemination events [41]. Further study will be needed to determine whether bradyzoites of this cyclin mutant have an enriched Group A transcriptome.

Overall, our findings revealed there are multiple subtypes of bradyzoites with distinct transcriptomes supporting the idea that tissue cysts are not homogenous and that a linear model of cyst maturation is unlikely. Furthermore, we identified novel bradyzoite markers that add to the current canonical markers and help distinguish natural bradyzoites from those generated in vitro. The data provides evidence as to why repeated culture and passage has led to vast variation in *Toxoplasma* strains used by the community. This puts the tissue cyst at the center of the *Toxoplasma* life cycle generating the flexibility required to maintain a chronic infection, initiate new infections and likely the adaptability to successfully invade multiple mammalian hosts. These findings open new avenues for investigating native bradyzoite biology and tissue cyst diversity.

## Methods and Materials

### Parasite maintenance and mouse infections

All animal experiments were approved by the Institutional Animal Care and Use Committee (IACUC) of the University of California, Riverside. Six-week-old female CBA, Swiss Webster (SWR/J) or C57Bl/6 mice were obtained from Jackson Laboratories (Jackson ImmunoResearch Laboratories, Inc., West Grove, PA, USA) and maintained in a pathogen-free vivarium. Type II ME49EW parasites were propagated *in vivo* by alternative passage between resistant (Swiss Webster, SWR/J) and sensitive (CBA/J) mouse strains, as previously described [41]. Mice were infected via intraperitoneal injection with 10 tissue cysts (in 200 μl) prepared from brain homogenates of mice infected for 30 days.

### Bradyzoite purification and ex vivo cultures following recrudescence

Following 40-day post-infection, ME49EW bradyzoites were isolated from infected mouse brains via Percoll gradient, as previously described [13]. Neonatal (from C57Bl/6 mouse pups) cortical astrocytes grown in T-75 cm2 vented flasks were used for *ex vivo* parasite cultures initiated by excysted bradyzoites. Primary astrocyte cultures were inoculated with excysted bradyzoites or day 3 and 5 parasite cultures at an MOI of 0.5 as previously described [13]. Infected cells were cultured in DMEM media supplemented with 5% heat-inactivated fetal bovine serum (FBS), 1% penicillin-streptomycin (Genclone), 1% GlutaMax supplement (Thermo Fisher Scientific, Waltham, MA, USA), 1% Na pyruvate and 2.5% HEPES buffer (Thermo Fisher Scientific Waltham, MA, USA) and placed in humidified hypoxia chambers containing 5 % oxygen and 5 % CO2 (nitrogen balanced).

### Production of antiserum against SRS22A

Rabbit antiserum was generated in-house against the *Toxoplasma* protein SRS22A using a selected peptide sequence (Fig. S1A, *LRGNDGRSSRVIEKEAEVAK*). Rabbits were immunized with the peptide, and antisera were collected at 28 days (first bleed), 56 days (second bleed), and 72 days (third bleed) post-immunization (Thermo Fisher Scientific Waltham, MA, USA).

### FACS sorting of bradyzoite populations

Upon excystation of purified ME49EW cysts, at least one million bradyzoites were stained for flow cytometry analysis using specific antibodies against SRS9 or SRS22A. Briefly, parasites were incubated with rabbit anti-SRS9 (kindly provided by John Boothroyd, Stanford University) or rabbit anti-SRS22A (generated in-house). Primary antibody incubation was performed at a 1:50 dilution in PBS in 5 ml round-bottom FACS tubes for 30 minutes at room temperature. Following incubation, each sample was washed with 1 ml of sterile PBS and centrifuged at 1500 rpm for 5 minutes. Secondary antibodies were diluted 1:1000 in PBS, with Alexa Fluor 647 goat anti-rabbit IgG (A-21244, Invitrogen, USA) used for SRS9 and Alexa Fluor 488 goat anti-rabbit IgG (A-11008, Invitrogen, USA) used for SRS22A. Parasites were then resuspended in 50 µl of diluted secondary antibodies and incubated for 30 minutes at room temperature in the dark. After incubation, stained parasites were washed with 1 ml of PBS, centrifuged, and resuspended in 300 µL of PBS for FACS analysis. Flow cytometry and cell sorting were performed using the MoFlo Astrios EQ Cell Sorter (Beckman Coulter) at the UC Riverside School of Medicine Flow Cytometry Core Facility to identify and sort distinct SRS9+ or SRS22A+ parasite populations. SRS9 and SRS22A expression were detected in the APC and FITC channels, respectively, and parasite populations were collected in 1 ml of infection media. Primary astrocyte cultures grown on coverslips were infected either with purified SRS22A+ or SRS22A− sorted parasites (500 parasites per well in 6-well plates, approximately MOI of 0.0004) or with SRS9+ parasites (2500 parasites per well in 24-well plates, approximately MOI of 0.01).

### Immunofluorescence staining

Population growth and SRS antigen expression patterns (SAG1 vs. SRS9) were assessed in primary astrocytes infected with sorted parasite populations on Day 5 post-infection. Cells were fixed in 4% paraformaldehyde (PFA), followed by three 1-minute washes with PBS. After permeabilization in acetone for 10 minutes, cells were washed three more times with PBS for 1 minute each. Next, cells were blocked with 5% donkey serum in PBS and incubated with antibodies specific for bradyzoites and tachyzoites, including rabbit anti-SRS9 (1:1000 dilution), mouse anti-SAG1 (1:1000 dilution, Bio-Rad, catalog # 9070-2020), and biotinylated DBA (1:500 dilution, B-1035-5, Vector Laboratories, CA, USA) to identify the cyst wall. Primary antibody incubation was carried out overnight at 4°C. The following day, cells were washed three times with PBS for 5 minutes each and incubated with secondary antibodies for 1 h at room temperature. The secondary antibodies used included donkey anti-rabbit Alexa Fluor 568 (1:1000 dilution, A10042, Invitrogen, USA), goat anti-mouse Alexa Fluor 488 (1:1000 dilution, A-11029, Invitrogen, USA), and Alexa Fluor 647-streptavidin conjugate (1:1000 dilution, S32357, Sigma-Aldrich, USA) to detect rabbit anti-SRS9, mouse anti-SAG1, and biotinylated DBA, respectively. After incubation, cells were washed three times with PBS for 10 minutes each and rinsed once with autoclaved water. DAPI VIBRANCE (Vectashield Vibrance, H-1800, Vector Laboratories, CA, USA) was then applied to each coverslip before mounting onto a glass slide. Parasite vacuoles were counted using a Leica DMI 6000 immunofluorescence microscope with a 40X objective, and vacuoles were enumerated.

### Single-cell RNA sequencing of in vivo bradyzoites

Single-cell RNA sequencing (scRNA-seq) was performed on excysted bradyzoites and parasite cultures obtained from ex vivo Day 7 astrocytes at a sequencing depth of one billion reads[13]. This was achieved using the 10X Genomics Chromium Next GEM Single Cell 3’ Reagent Kit v3.1, as previously described [13]. Sequencing was conducted at the San Diego IGM Genomics Center using an Illumina NovaSeq 6000. As before, Cell Ranger version 5.0.1 was used for read alignment to a custom reference genome (ME49 *Toxoplasma gondii* reference genome) and for count matrix generation. scRNA-seq data was analyzed using the Seurat package (R version 4.4.2 and Seurat 5.1.0) with libraries ‘ggplot2’, ‘tidyverse’, ‘conflicted’, ‘gridExtra’, ‘dplyr’, ‘EnhancedVolcano’ and ‘ggrepel’. Dimensionality reduction was first performed using Principal Component Analysis (PCA), followed by construction of a Shared Nearest Neighbor (SNN) graph via the ‘FindNeighbors’ function. Cells were then clustered using the Louvain algorithm at a resolution of 0.2, yielding five transcriptionally distinct clusters. For visualization, cells were embedded in two dimensions using Uniform Manifold Approximation and Projection (UMAP).’ A non-parametric Wilcoxon rank test was applied to identify differentially expressed genes among Groups A-E. Volcano plots were generated from a comparison of each bradyzoite Group A-E (chronic phase infection) and the BAG1+ bradyzoites (early phase) that spontaneously arise in our ex vivo bradyzoite recrudescence model at Day-7 as explained in Results.

### Bradyzoite differentiation assay under alkaline stress

Alkaline stress assay was performed in HFFs as previously described [51, 52]. A bradyzoite differentiation media was prepared using RPMI 1640 (GIBCO) medium supplemented with 5% FBS, 50 mM HEPES, 100 unit/ml penicillin, and 100 μg/ml streptomycin. The alkaline medium pH was adjusted to 8.2 using NaOH. Briefly, HFFs were infected with ME49B7 parasites for 4 h, and then media was removed and rinsed four times with parasite media (5% FBS, 1% Gln, 100 unit/ml penicillin, and 100 μg/ml streptomycin) to remove unattached parasites. The bradyzoite differentiation media was added onto the cells and cell culture flasks were placed at 37C for 48 h. Cells were fixed in 4% PFA post-infection and then stained for SAG1, SRS9 and SRS22A using immunofluorescence staining as described above.

### Preparation of PECs and brain homogenates at days 5- and 28- post-infection

On 5-days post-infection, mice were anesthetized and injected intraperitoneally with 5 ml of cold PBS with 1% BSA. Peritoneal exudate cells were collected by withdrawing the injected solution and subsequently enumerated. For each mouse, 100,000 cells were deposited onto microscope slides using cytospin funnels and centrifugation at 1200 rpm for 5 minutes. On 28-days post-infection, infected mouse brains were harvested in 3 ml cold PBS, and brain homogenates were prepared by sequentially passing the tissue through 18G, 20G, and 22G needles. A 30 ul aliquot of brain homogenate was deposited on a microscope slide, smeared using a coverslip and allowed to dry. Slides were then fixed in methanol and stored at -20°C until immunofluorescence staining was performed.

### Detection of parasite burden in mouse tissues

Liver, spleen and lung were harvested from infected mice and stored at -80°C prior to DNA purification. DNA was purified using a DNeasy minikit (Qiagen) according to manufacturer’s instructions. The qPCR reaction used 600 ng of DNA from each organ, 0.375 μM B1 gene primers (Forward:5′- TCCCCTCTGCTGGCGAAAAGT-3′; Reverse: 5′-AGCGTTCGTGGTCAACTATCG-3′), and 2x SensiFAST SYBR No-ROX qPCR master mix (Thomas Scientific, #C755J00) in a 20 μl total volume. The qPCR program was: 3 min at 95°C, 40 cycles of 10s at 95°C for denaturation, 30s at 60°C for annealing, followed by 5s at 65°C and 5s at 95°C for extension. The real-time data was collected on a BioRad CFX96 Touch Real-Time PCR Detection system, using Bio-Rad CFX software. A standard curve was generated with serial dilution of purified ME49 genomic DNA and used to calculate pg amount from the equation displayed on the linear trendline (R^2^=0.99). All results were expressed as the pg of parasite genomic DNA calculated from an experimentally generated standard curve and relative expression was presented on each graph as compared to naïve controls.

### Western blot analysis of SRS22A

Cysts were purified from the brains of three CBA mice infected with ME49EW for 30 days. Bradyzoites were excysted from the pooled cysts by 90 seconds of pepsin-HCl treatment at 37C. RH grown in HFFs were used as a negative control. These parasites were lysed by a 25G needle and filtered in a 3µm filter. After washing the parasites, protein lysates were made by adding 100μL NP-40 buffer with protease inhibitors for each 1×10⁶ pelleted parasites. After keeping on ice for 10 minutes, the lysate was centrifuged at 14,000xg for 10 minutes at 4°C, and the supernatant collected. 1μg of each protein lysate was separated by SDS-PAGE (15% resolving gel, 5% stacking gel) for 2 h and transferred onto a 0.2µm nitrocellulose membrane for 35 minutes. Membranes were incubated in 5% milk in 0.1% PBS-T for 30 minutes at room temperature, then probed with rabbit-α-SRS22A antibody (1:500) or pre-immune serum (1:500) overnight at 4°C. After washing, membranes were probed with goat-α-rabbit HRP conjugate (Bio-Rad #1705046, 1:10,000) for one h at room temperature. The membranes were then reprobed with rabbit-α-PCNA antibody (1:2,500) overnight at 4°C, followed by goat-α-rabbit HRP conjugate for one h at room temperature. Importantly, signal was detected with the SuperSignal™ West Femto Maximum Sensitivity Substrate (ThermoScientific #34094). Membranes were imaged on the Bio-Rad Gel Doc XR+ System at an exposure of 5 seconds.

### Statistical analysis

Statistical analyses for tissue cyst diameter, cyst counts, SRSR22A staining intensity, vacuole counts, and parasite burden were conducted in GraphPad Prism (version 10.2.3) using one-way or two-way ANOVA or two-tailed t-tests, as appropriate. A P-value <0.05 was considered statistically significant. scRNA-seq data were analyzed using Seurat [53] as described above.

## Supporting information

Database S1

Database S2

## Data and code availability

scRNA-seq data were deposited into GEO **(accession number).** No custom code was generated for scRNA-seq analysis in this study. Analysis pipelines used are available from the authors upon request.

## Acknowledgements

*Toxoplasma gondii* gene information was accessed on http://ToxoDB.org. We thank Dr. John Boothroyd for the antibodies used in our study. We would like to acknowledge Mary Hamer, who is the manager of Research Core at UC Riverside School of Medicine for her help with FACS cell sorting operations. We also acknowledge Dr. Brandon Le from the UC Riverside IIGB Bioinformatics Core for his help with the analysis of the scRNAseq data. This study was supported by NIH NIAID R01 grants AI158417 and AI124682 to M.W.W. and E.H.W., AI122760 to M.W.W., and DA048815 to E.H.W.

## Supplementary Figure Legends

**Figure S1. [A.]** Peptide selected for generating a monoclonal SRS22A antibody in rabbit. **[B.]** Immunofluorescence images of co-staining of ME49EW cyst using rabbit-a-SRS22A antibody (green), pre-immune control (green) and DBA (red). **[C.]** Examples of intermediate staining of parasite vacuoles (SAG1-green and SRS9-red) in FACS sorted SRS22A negative infections in astrocytes 5-days post-infection after FACS sorting (top panel), and examples of secondary infections in both FACS sorted SRS22A + and – infections in astrocytes on 5-days post-infection following FACS sorting.

**Figure S2. [A. and B.]** Compensation controls used in FACS purification of SRS22A positive and negative populations from 40-day infected mice and the purity of sorted populations (A) and (B) SRS9 positive and negative populations from ex vivo Day 7 cultures from astrocytes infected with ex vivo bradyzoites.

**Figure S3. [A.]** Number of unique molecular identifiers (UMIs) per cell is shown for each sample included in the analysis. A 300 UMI threshold cutoff was applied to all samples to filter out poor quality parasites. We also performed a correlation analysis between the two in vivo bradyzoite samples (excysted bradyzoites from 40-day infected mice tissue cysts) included in the scRNAseq analysis. A strong correlation was observed, Pearson correlation coefficient of r=0.965. **[B.]** Quality assurance parameters of scRNA seq are shown. **[C.]** UMAP projections of bradyzoites from two independent harvests of 40-day brain tissue cysts show identical clustering. **[D.]** Table of selected growth gene expression (average expressed values) across Group A-E bradyzoites. Note that growth gene expression was uniformly higher in Group E bradyzoites, which constitutes 4% of the total bradyzoites analyzed in this experiment (4,181 total bradyzoites). The proportional value of the other bradyzoite groups is also indicated. **[E.]** The elbow plot indicates a noticeable decrease in variation after principal component 18. Pearson correlations between cell embeddings from PCs 1-18 PCs and cell scores for G1, S/M and bradyzoite specific gene sets are shown. **[F.]** UMAP projection of Groups A-E as compared to the BAG1+ bradyzoites from ex vivo bradyzoite-infected astrocytes at day 7. A distinct clustering of Groups A-E from the D7-BAG1+ parasites is easily seen. The Table below shows the total number of down and upregulated genes as well as the unique upregulated genes in each Group A-E. **[G.]** UMAP projection displaying SRS22A expression shading shows a SRS22A expression is enhanced in Group B bradyzoites, while absent from D7-BAG1+ bradyzoites.

**Figure S4. [A.] (top row)** UMAP projection of in vivo bradyzoites references Groups A-E distributions. Violin plots of LDH2 and AP2IX-9 are shown for Groups A-E. (**second row**) UMAP mRNA shading of bradyzoite-specific rhoptry protein, BRP1, and associated violin plot are shown. The last panel shows an example of unique rhoptry gene expression (TGME49_308093) in Group A bradyzoites. (**third row**) UMAP projection of GRA82 mRNA levels showing enhanced expression in Groups A-C (3-7-fold higher than Groups D & E, Database S1), and UMAP images of SRS9 mRNA and cyst wall protein, SRS13 mRNA enhanced expression in Groups C and D bradyzoites. A heatmap of GRA **[B.]** and ROP/RON **[C.]** family gene expression across Groups A-E bradyzoites. The right heat map in (C) shows the analysis of representative S/M transcripts in Group A versus E bradyzoites. Note the increase of rhoptry mRNA expression in Group E is accompanied by increased expression of other S/M transcripts, while enhanced rhoptry mRNA expression of Group A bradyzoites did not show this cell cycle pattern.

## Notes

### Competing Interest Statement

The authors have declared no competing interest.

