## Supplementary figures and images for "Bradyzoite subtypes rule the crossroads of *Toxoplasma* development"

### Supplemental Figure S1

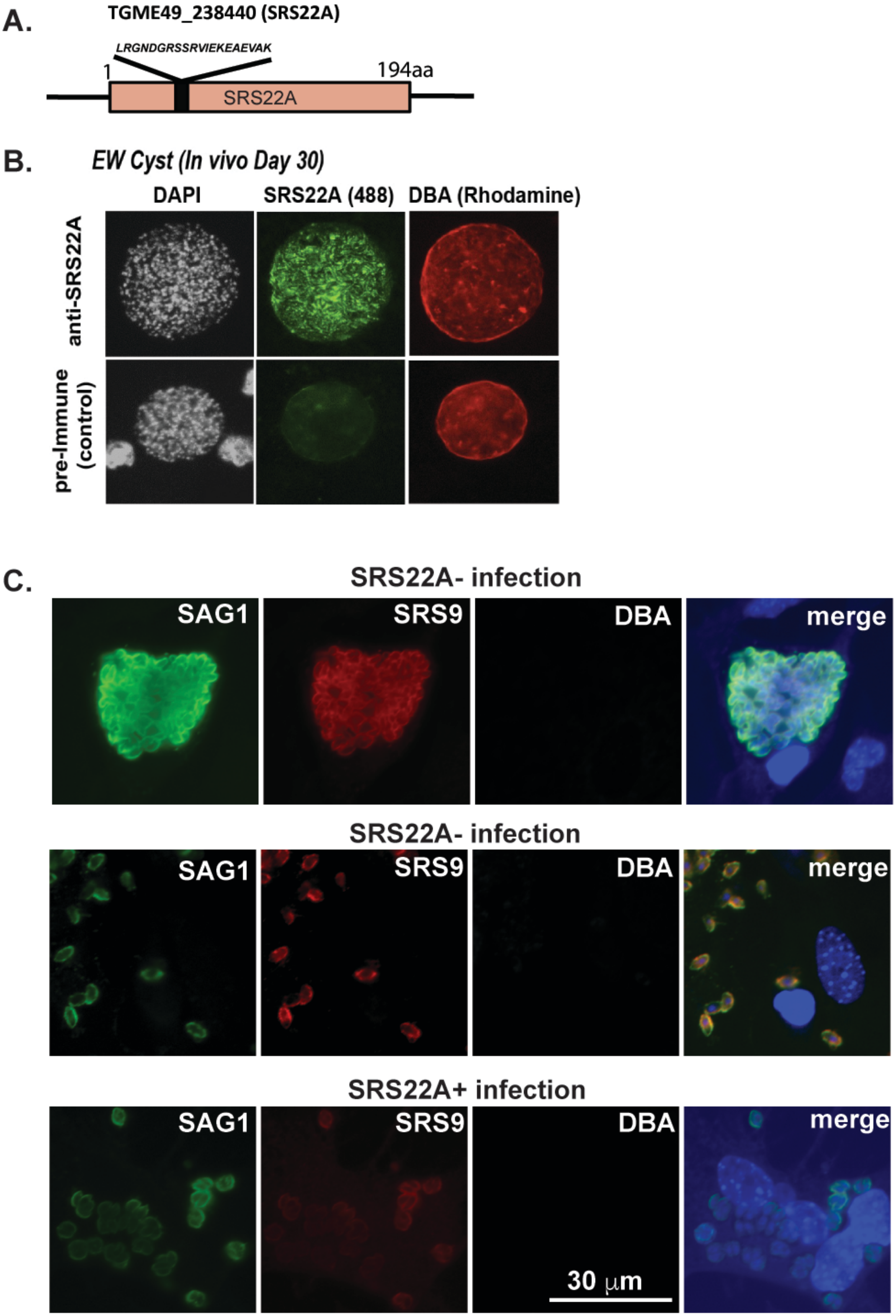

### Supplemental Figure S2

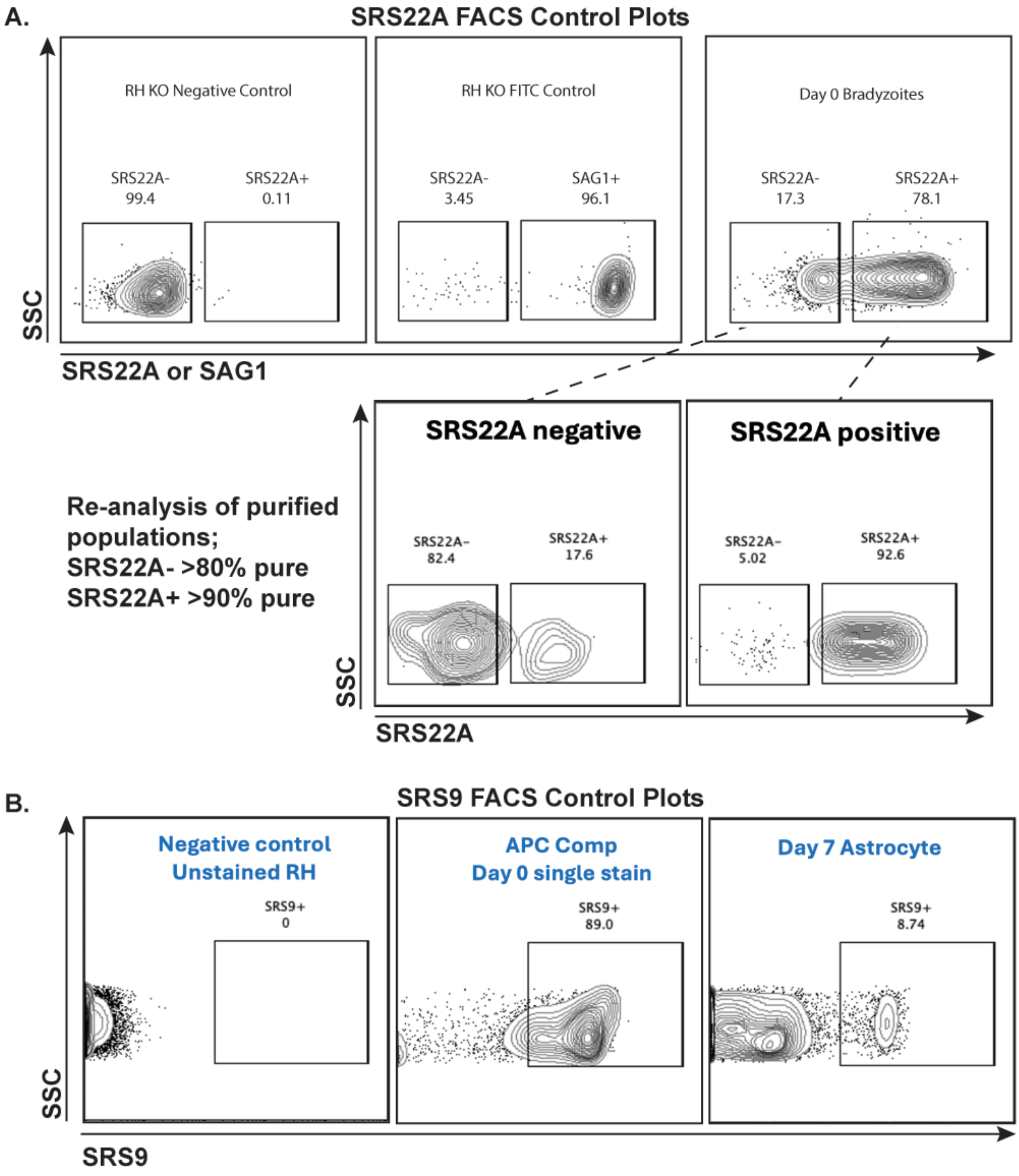

### Supplemental Figure S4

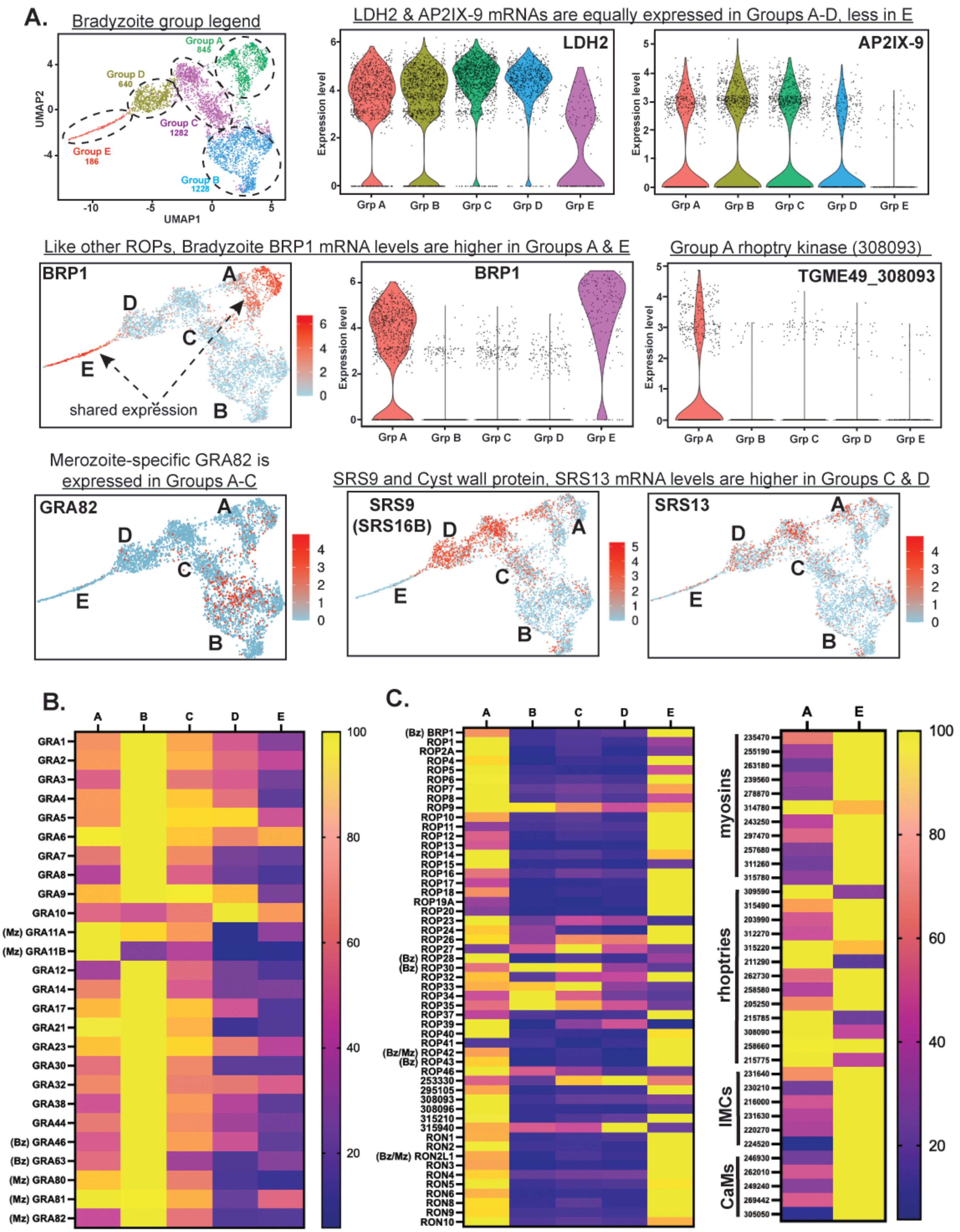
