## Supplemental Figure S3 for "Bradyzoite subtypes rule the crossroads of *Toxoplasma* development"

A.

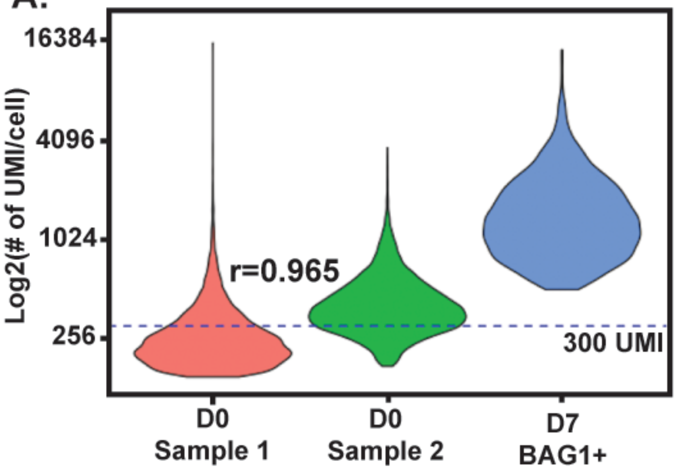


B.

| Library | D0-1 | D0-2 | Day-7 Astro |
|---|---|---|---|
| Total reads (x10^8) | 7.24 | 15.4 | 8.76 |
| Mapped reads (fraction reads in cells) | 81.40% | 82.50% | 80.60% |
| # Cells | 8,609 | 9,153 | 29,931 |
| Mean reads/cell | 84,205 | 168,724 | 32,523 |
| Median reads/cell | 177 | 267 | 511 |
| Total genes detected | 7,705 | 7,978 | 8,115 |
| > 300 UMI cell counts | 888 | 3,302 | 26,414 |

C.

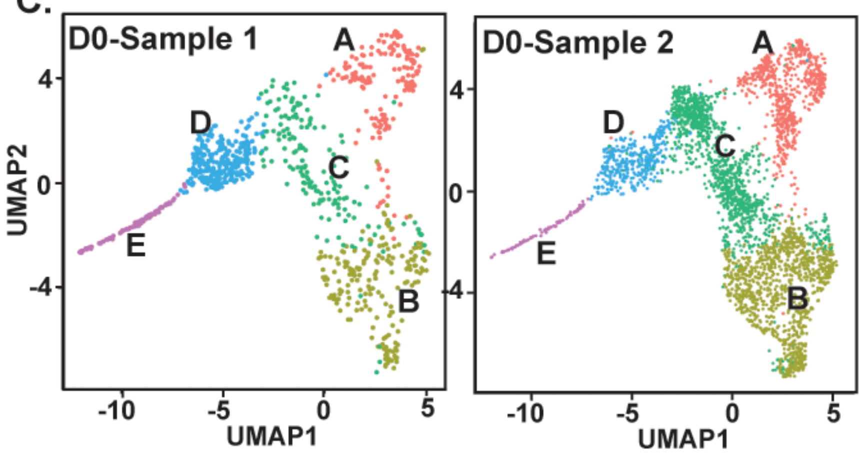


F.

Comparison of early in vitro bradyzoites (D7-BAG1+) to in vivo bradyzoites (A-E)

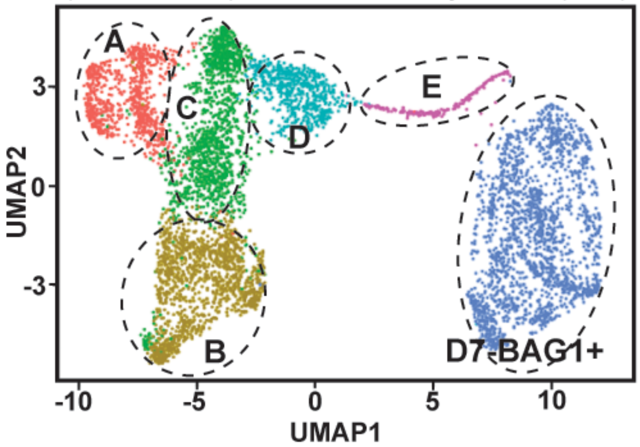


Gene numbers per bradyzoite Group A-E compared to early D7-BAG1+ bradyzoites

| Group | #up-reg | #down-reg | #unique up-reg |
|---|---|---|---|
| A | 401 | 243 | 109 |
| B | 385 | 543 | 103 |
| C | 298 | 264 | 67 |
| D | 359 | 278 | 32 |
| E | 495 | 259 | 173 |

D.

Selected growth gene comparison.
GRA1 and GRA2 are consitutive controls

| Gene ID TgME49_ | Gene name | Grp A **20% | Grp B 30% | Grp C 31% | Grp D 15% | Grp E 4% |
|---|---|---|---|---|---|---|
| 270250 | GRA1 | 3.4 | 4.8 | 3.7 | 2.5 | 1.4 |
| 227620 | GRA2 | 2.8 | 3.9 | 3.1 | 2.2 | 1.7 |
| 209910 | H2B | 2.2 | 1.1 | 1.5 | 1.8 | 3.6 |
| 300200 | H2A | 1.5 | 1.2 | 1.3 | 1.4 | 3.0 |
| 316400 | aTUB | 1.4 | 0.4 | 0.7 | 1.5 | 3.8 |
| 266960 | bTUB | 1.2 | 0.6 | 1.0 | 2.3 | 3.4 |
| 250340 | Centrin 2 | 0.2 | 0.07 | 0.01 | 0.2 | 2.0 |
| 260820 | ISP1 | 1.1 | 0.3 | 0.3 | 0.4 | 2.8 |
| 231640 | IMC1 | 1.5 | 0.6 | 0.6 | 0.3 | 2.1 |
| 216000 | IMC3 | 1.2 | 0.2 | 0.2 | 0.09 | 2.4 |

**% of total bradyzoites sequenced and >300 UMI (4,181 total Bz).

G.

SRS22A mRNA expression

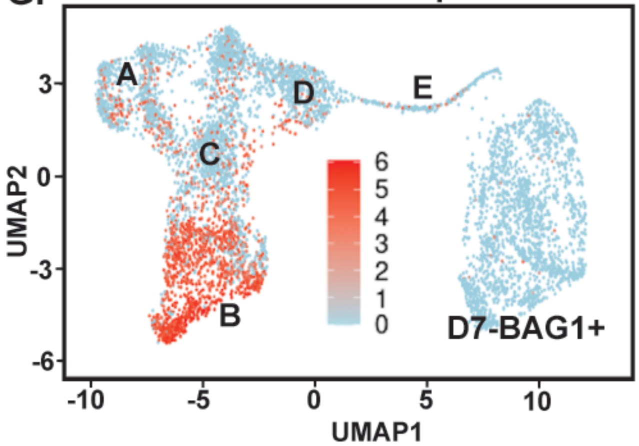


E.

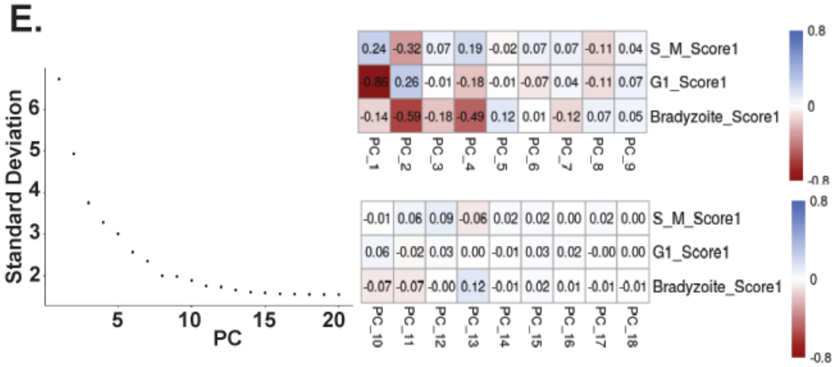
